# MEN1 deficiency creates a selenite-dependent switch in ferroptosis

**DOI:** 10.64898/2026.07.16.738891

**Authors:** Ronnie Ren Jie Low, Nishita Parnandi, Aurora I Idilli, Ersa Gjelaj, Fernanda T. Subtil, Sandra Segura-Bayona, James I. MacRae, Simon J. Boulton

**Affiliations:** DSB Repair Metabolism Laboratory, The Francis Crick Institute; London NW1 1AT, UK; Metabolomics STP, The Francis Crick Institute; London NW1 1AT, UK

## Abstract

Replication stress is a hallmark of cancer cells, yet the factors determining tolerance remain poorly understood. Genome-wide CRISPR screens for modifiers of replication-stress responses identified loss of the chromatin scaffold gene MEN1 as a resistance factor. We show that MEN1 deficiency suppresses lipid peroxidation and ferroptotic death, enabling survival following treatment with multiple replication stress-inducing agents. Mechanistically, MEN1 loss reduced H3.3 occupancy at ACSL1 regulatory regions and lowered ACSL1 expression, with ACSL1 loss phenocopying replication stress resistance. MEN1- or ACSL1-deficient cells also exhibited reduced levels of SLC7A11 and glutathione, rendering them hypersensitive to GPX4 inhibition under selenite-limiting conditions. Conversely, selenite supplementation preferentially increased GPX4 abundance and converted these cells to a ferroptosis-resistant state independently of SLC7A11. Thus, MEN1 links chromatin regulation to the ferroptotic control of replication stress responses, while selenite availability determines whether MEN1-deficient cells are vulnerable or resistant.

## Introduction

DNA replication is continually challenged by endogenous DNA lesions, nucleotide depletion, difficult-to-replicate sequences and oncogene activation, as well as by chemotherapeutic agents that impede DNA synthesis. These challenges generate replication stress and activate checkpoint, fork-protection and DNA-repair pathways that preserve genome stability and cell viability^1^. When these responses are overwhelmed, stalled replication forks collapse, oxidative stress increases and cells undergo irreversible arrest or death^1,2^. The outcome of replication stress therefore depends not only on lesion processing at stalled forks, but also on the pathways that execute or suppress cell death. Although apoptosis and mitotic catastrophe are established consequences of replication stress^3^, the contribution of other regulated death pathways remains less well understood. Whether ferroptosis determines cellular tolerance to replication stress, and which genetic factors connect replicative damage to lipid peroxidation, remain unclear.

Ferroptosis is an iron-dependent form of regulated cell death driven by the peroxidation of polyunsaturated fatty acid-containing phospholipids^4^. Ferroptotic sensitivity is determined by the balance between the generation of peroxidation-prone lipids and antioxidant systems that detoxify phospholipid hydroperoxides. ACSL4 has an established role in activating polyunsaturated fatty acids for incorporation into membrane phospholipids, whereas the related enzyme ACSL1 displays context-dependent pro- and anti-ferroptotic activities ^5–9^. The principal antioxidant defence is the SLC7A11-glutathione-GPX4 axis, which couples cystine uptake and glutathione synthesis to GPX4-dependent reduction of phospholipid hydroperoxides, alongside FSP1, which provides a parallel protective pathway^10–12^. As a selenoprotein, GPX4 synthesis depends on extracellular selenite availability, which dictates cellular resistance to ferroptotic stress^13^. Accordingly, differences in the selenite content of culture media or serum can markedly alter GPX4 abundance and ferroptosis sensitivity^14^. Thus, genetically encoded ferroptosis phenotypes may be revealed, masked or even reversed by nutrient availability. How such environmental inputs intersect with chromatin-dependent transcriptional programmes to determine ferroptotic cell fate remains poorly understood.

MEN1 encodes menin, a nuclear scaffold that coordinates transcriptional and chromatin-regulatory complexes^15^. Its function is highly context dependent: menin acts as an oncogenic cofactor in KMT2A-rearranged leukaemias, whereas MEN1 loss promotes tumorigenesis in endocrine tissues^16^. Pancreatic neuroendocrine tumours frequently harbour MEN1 loss-of-function mutations, which often coexist with alterations in ATRX or DAXX^17^. The histone variant H3.3 is deposited at gene regulatory regions by the HIRA chaperone complex and at repetitive regions, including telomeres, by ATRX-DAXX complex^18–20^. As a consequence, loss of ATRX or DAXX is strongly associated with alternative lengthening of telomeres^21^. Recently, ATRX-deficient cells have been reported to show increased replication stress and ferroptosis sensitivity^21,22^.

Here, analysis of genome-wide CRISPR-Cas9 loss-of-function screens performed in independent cellular models exposed to replication stress-inducing agents identified MEN1 loss as a recurrent resistance factor. MEN1 deficiency conferred resistance to replication stress inducing agents and suppressed ferroptotic death. Our investigation of the underlying resistance mechanism revealed that MEN1 loss reduced H3.3 occupancy at *ACSL1* regulatory regions and lowered ACSL1 expression. Similarly, HIRA deficiency reduced ACSL1 expression, and ACSL1 loss phenocopied replication stress resistance. Unexpectedly, MEN1- or ACSL1-deficient cells also exhibited reduced SLC7A11 abundance and diminished glutathione levels, creating an intrinsic sensitivity to GPX4 inhibition under selenite-limiting conditions. Conversely, selenite supplementation preferentially increased GPX4 abundance and converted these cells to a ferroptosis-resistant state independently of SLC7A11. Together, these findings connect MEN1-dependent chromatin regulation to lipid and redox metabolism and show that selenite availability can reverse the cell death phenotype associated with MEN1 loss. More broadly, they establish metabolic control of ferroptosis as a determinant of cellular tolerance to replication stress.

### MEN1 deficiency confers ferroptosis resistance

Many chemotherapeutic agents induce replication stress, elevating ROS and DNA damage, and ultimately triggering cell death^2,23^. MEN1 deficiency was found to confer resistance to replication stress-inducing drugs hydroxyurea (HU) and camptothecin (CPT) in genome-wide CRISPR-Cas9 loss-of-function (LOF) screens using two different cellular models: eHAP-iCas9 and RPE1-hTERT Cas9 TP53^-/-^ (Fig. 1a and Extended Data Fig. 1a) ^21,24^.

**Fig. 1.**
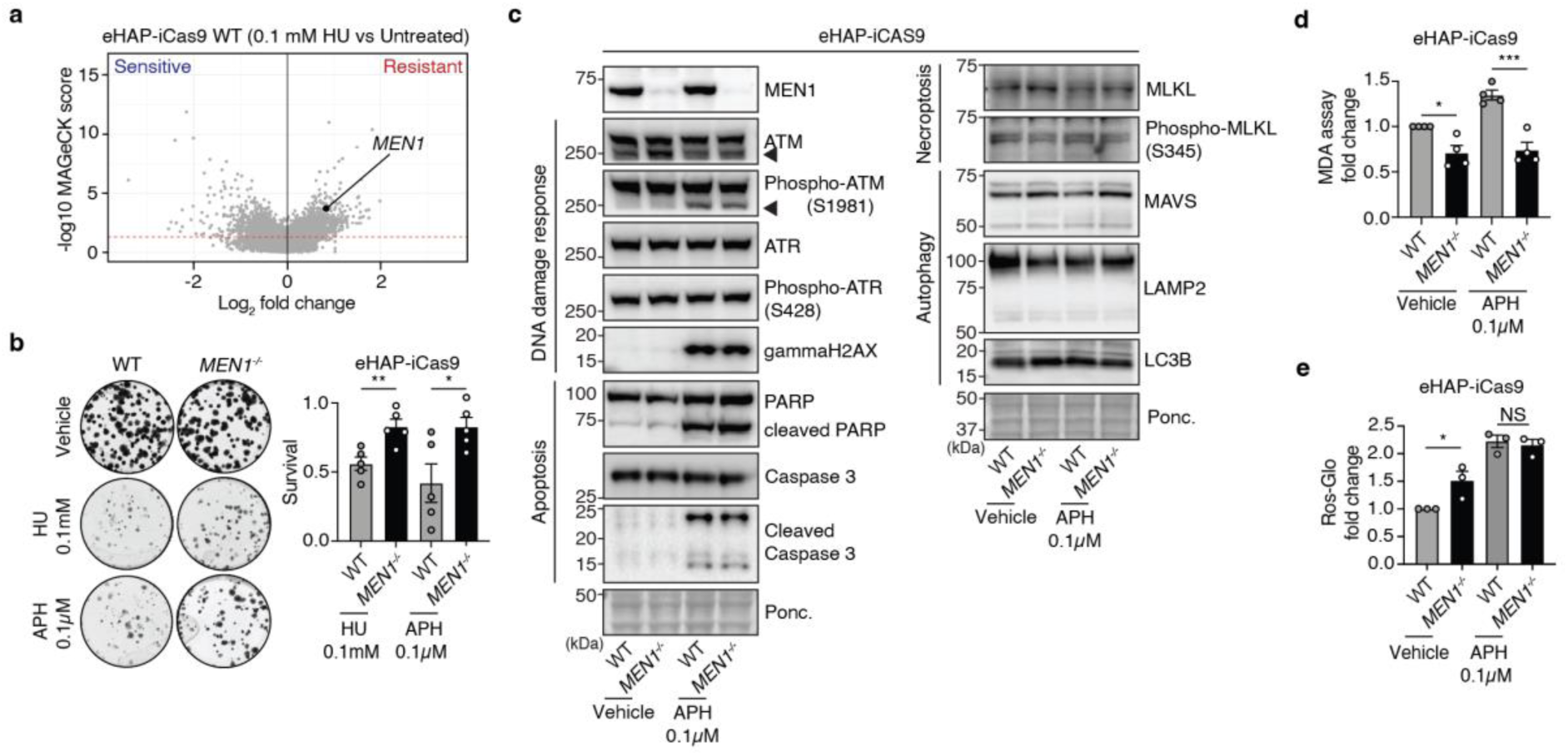
MEN1 deficiency suppresses ferroptosis to confer resistance to replication stress. **a**, Volcano plot of a genome-wide CRISPR-Cas9 LOF synthetic lethal screen in eHAP-iCas9 WT cells treated with 0.1 mM hydroxyurea (HU) versus untreated controls. *MEN1* is highlighted as a significant resistance hit. Dashed red line indicates the significance threshold (MAGeCK score = 1)^21^. **b**, Representative colony formation assay (CFA) images (left) and quantification (right) of WT and *MEN1*^-/-^ eHAP-iCas9 cells treated with vehicle, 0.1 mM HU, or 0.1 μM aphidicolin (APH) for 5 days. Colony counts are normalised to WT treated with vehicle. *MEN1*^-/-^ cells show significantly increased survival under both replication stress agents (**p < 0.01, *p < 0.05; Student’s t test). Mean ± SEM; n =5 independent experiments. **c**, Immunoblot analysis of DDR pathway markers and cell death pathways in WT and *MEN1*^-/-^eHAP-iCas9 cells with or without APH treatment. No differential expression in DDR markers (phospho-ATM, phospho-ATR, and γ-H2AX), apoptosis markers (PARP, Caspase 3), necroptosis markers (MLKL), and autophagy markers (MAVS, LAMP2, LC3B) is observed in *MEN1*^-/-^ cells. **d**, Quantification of malondialdehyde (MDA) levels, a marker of lipid peroxidation, by MDA assay in WT and *MEN1*^-/-^ eHAP-iCas9 cells treated with vehicle or 0.1 μM APH. Values are normalised to WT treated with vehicle. MDA is significantly reduced in *MEN1*^-/-^ cells relative to WT under both vehicle and APH conditions, indicating suppression of ferroptosis-associated lipid peroxidation (*p < 0.05, ***p < 0.001; one-way ANOVA with Tukey’s post hoc test). Mean ± SEM; n =4 independent experiments. **e**, Quantification of reactive oxygen species (ROS) levels measured by ROS-Glo assay in WT and *MEN1*^-/-^ eHAP-iCas9 cells treated with vehicle or 0.1 μM APH for 24 hours. Values are normalised to WT treated with vehicle. Both WT and *MEN1*^-/-^ cells show elevated ROS upon APH treatment, confirming replication stress-induced oxidative burden. Mean ± SEM; n =3 independent experiments.

To validate these findings, we generated *MEN1*^-/-^ eHAP-iCas9 cells (Fig. 1c) and observed increased resistance following treatment with HU, aphidicolin (APH; a DNA polymerase-α inhibitor) and CPT in cell viability assays (Fig. 1b and Extended Data Fig. 1b). To explore the mechanistic basis of this resistance, we first evaluated different replication stress-associated cell death pathways^25^. We excluded roles for MEN1 in DNA damage response activation, apoptosis, necroptosis and autophagy, which might otherwise have explained the replication stress resistance of MEN1-deficient cells (Fig. 1c and Extended Data Fig. 1c). Instead, *MEN1*^-/-^ cells exhibited reduced levels of malondialdehyde, a marker of lipid peroxidation and ferroptosis upon APH treatment when compared to wild-type (WT) cells (Fig. 1d). Notably, APH treatment increased ROS in both WT and *MEN1*^-/-^ cells (Fig. 1e) indicating that MEN1 deficiency confers resistance to replication stress-induced cell death downstream of ROS production.

*As MEN1* mutations frequently co-occur with loss-of-function alterations in the H3.3 histone chaperones *ATRX* or *DAXX* in pancreatic neuroendocrine tumours (Fig. 2a) and given that ATRX-deficient cells are more susceptible to replication stress and ferroptosis sensitivity^21,22^, we examined whether MEN1 deficiency modifies ferroptosis in these contexts. Genome-wide CRISPR-Cas9 LOF screens in *ATRX*^-/-^ eHAP-iCas9 cells treated with HU again identified *MEN1* as a resistance hit (Fig. 2b)^21^. We validated this by generating MEN1 knockout cells in WT, *ATRX*^-/-^ and *DAXX*^-/-^ backgrounds (Fig. 2c). MEN1 deficiency conferred resistance to HU and APH treatment across all tested genetic backgrounds (Fig. 2d and Extended Data Fig. 2a,b).

**Fig. 2.**
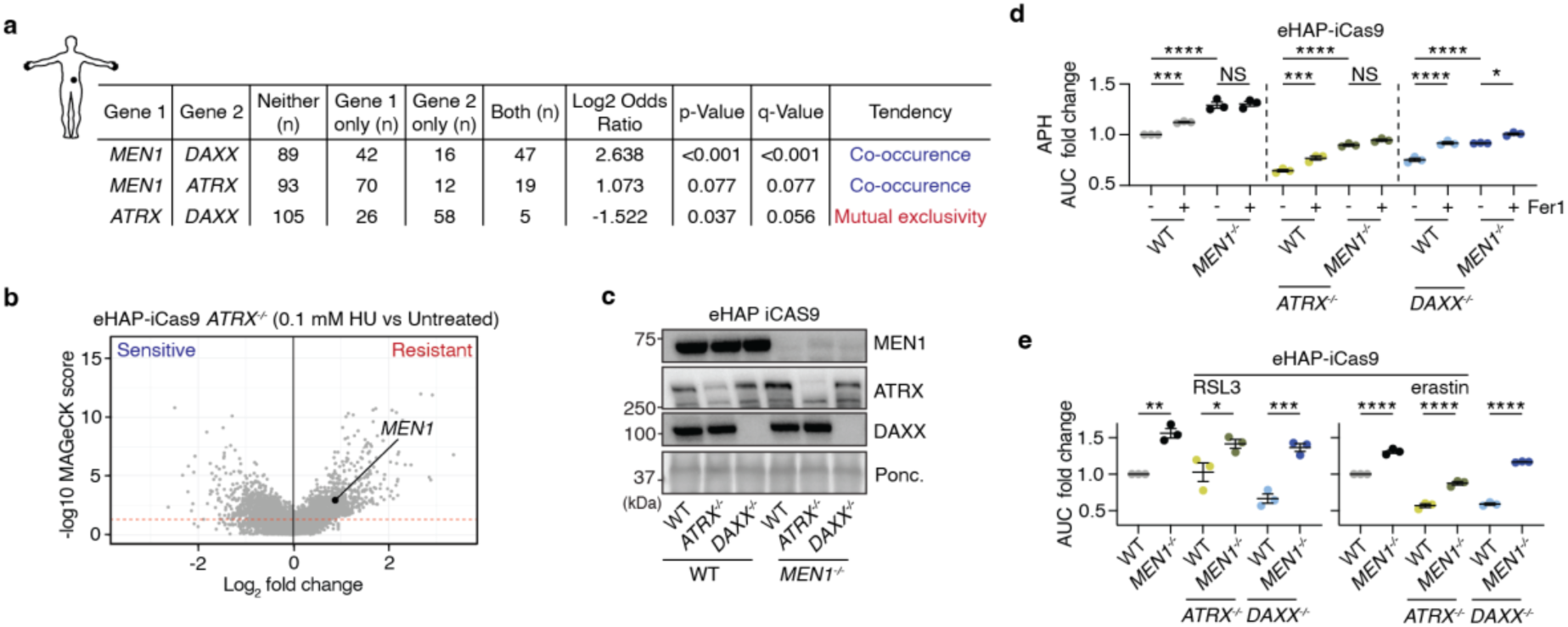
MEN1 deficiency confers ferroptosis resistance in ATRX- and DAXX-deficient backgrounds. **a**, Co-occurrence analysis of somatic mutations in *MEN1*, *ATRX*, and *DAXX* in pancreatic neuroendocrine tumour (PNET) patient data from cBioPortal. P value (Fisher’s exact test), and q value (Benjamini-Hochberg correction) are shown. *MEN1* and *ATRX* or *DAXX* mutations co-occur; *ATRX* and *DAXX* mutations are mutually exclusive. **b**, Volcano plot of a genome-wide CRISPR-Cas9 LOF synthetic lethal screen in *ATRX*^-/-^ eHAP-iCas9 cells treated with 0.1 mM HU versus untreated controls^21^. **c**, Immunoblot validating loss of MEN1, ATRX, and DAXX protein in the indicated eHAP-iCas9 single- and double-knockout cell lines. Cells were generated by CRISPR-Cas9 targeting using the indicated crRNAs (crNTC control, *MEN1*^-/-^) in WT, *ATRX*^-/-^, or *DAXX*^-/-^ backgrounds. Ponceau S staining serves as loading control. **d**,**e** Area under the curve (AUC) quantification of cell viability dose-response assays in the indicated genotypes (WT and *MEN1*^-/-^ in WT *ATRX*^-/-^, or *DAXX*^-/-^ backgrounds) from Extended Data Fig. 2a,b,c. APH dose-response ± 20 μM ferrostatin-1 (Fer-1), a ferroptosis inhibitor. Fer-1 significantly rescues APH-induced cell death in WT but not *MEN1*^-/-^ cells (**d**). MEN1 loss additionally confers ferroptosis resistance in *ATRX*^-/-^ and *DAXX*^-/-^ backgrounds with RSL3, a GPX4 inhibitor, and Erastin, a SLC7A11 inhibitor (**e**). (*p < 0.05, **p < 0.01, ***p < 0.001, ****p < 0.0001; one-way ANOVA with Tukey’s post hoc test). Mean ± SEM; n =3 independent experiments.

Consistent with a role for MEN1 in regulating ferroptosis, ferrostatin-1 (Fer1), a ferroptosis inhibitor, rescued APH-induced cell death in all tested genetic backgrounds but did not further enhance resistance in cells deficient for MEN1 (Fig. 2d and Extended Data Fig. 2b). We also tested ferroptosis inducers targeting the SLC7A11-GSH-GPX4 axis, including the GPX4 inhibitor, RSL3, and the SLC7A11 inhibitor, erastin, in WT *ATRX*^-/-^ and *DAXX*^-/-^ backgrounds ±MEN1 (Fig. 2e and Extended Data Fig. 2c). MEN1-deficiency reduced sensitivity to these agents in all genetic backgrounds, indicating that MEN1 loss suppresses ferroptotic cell death under replication stress and following ferroptosis induction in WT or ATRX/DAXX deficient backgrounds.

Although MEN1 overexpression has been reported to promote ferroptosis via the mTOR-SREBP1-SCD1 axis in PNET cell line models^26^, we found no evidence of enhanced SREBP1 processing or SCD1 upregulation in MEN1-deficient eHAP-iCas9 or in the human pancreatic neuroendocrine tumour (PNET) cell line, BON-1. Instead, SCD1 was reduced, likely as a secondary adaptation to altered lipid balance (Extended Data Fig. 2d). Similarly, a study in small cell lung carcinoma linked MEN1 deficiency to ferroptosis suppression via CD44v6 and COX2 downregulation^27^. However, we observed no changes in these proteins in our cell models (Extended Data Fig. 2d), potentially reflecting context- or tissue-specific differences.

### MEN1 deficiency affects ACSL1 expression through the HIRA-H3.3 axis

Loss of ATRX or DAXX has also been linked to altered tolerance of replication stress-induced cell death, raising the possibility that shared H3.3-dependent mechanisms could modulate ferroptosis sensitivity. Because MEN1 itself has been linked to chromatin regulation and genome stability^15^, we asked whether MEN1 influences H3.3 dynamics (Fig. 3a). H3.3 CUT&Tag sequencing revealed both genome-wide and telomeric alterations in H3.3 occupancy in MEN1^-/-^cells (Fig. 3b and Extended Data Fig. 3a). Co-mutation of *MEN1* with *ATRX* or *DAXX* maintained ALT-associated features (ss-TeloC and C-circles), whereas MEN1 deficiency alone produced more limited effects (Fig. 3c and Extended Data Fig. 3b)^17^.

**Fig. 3.**
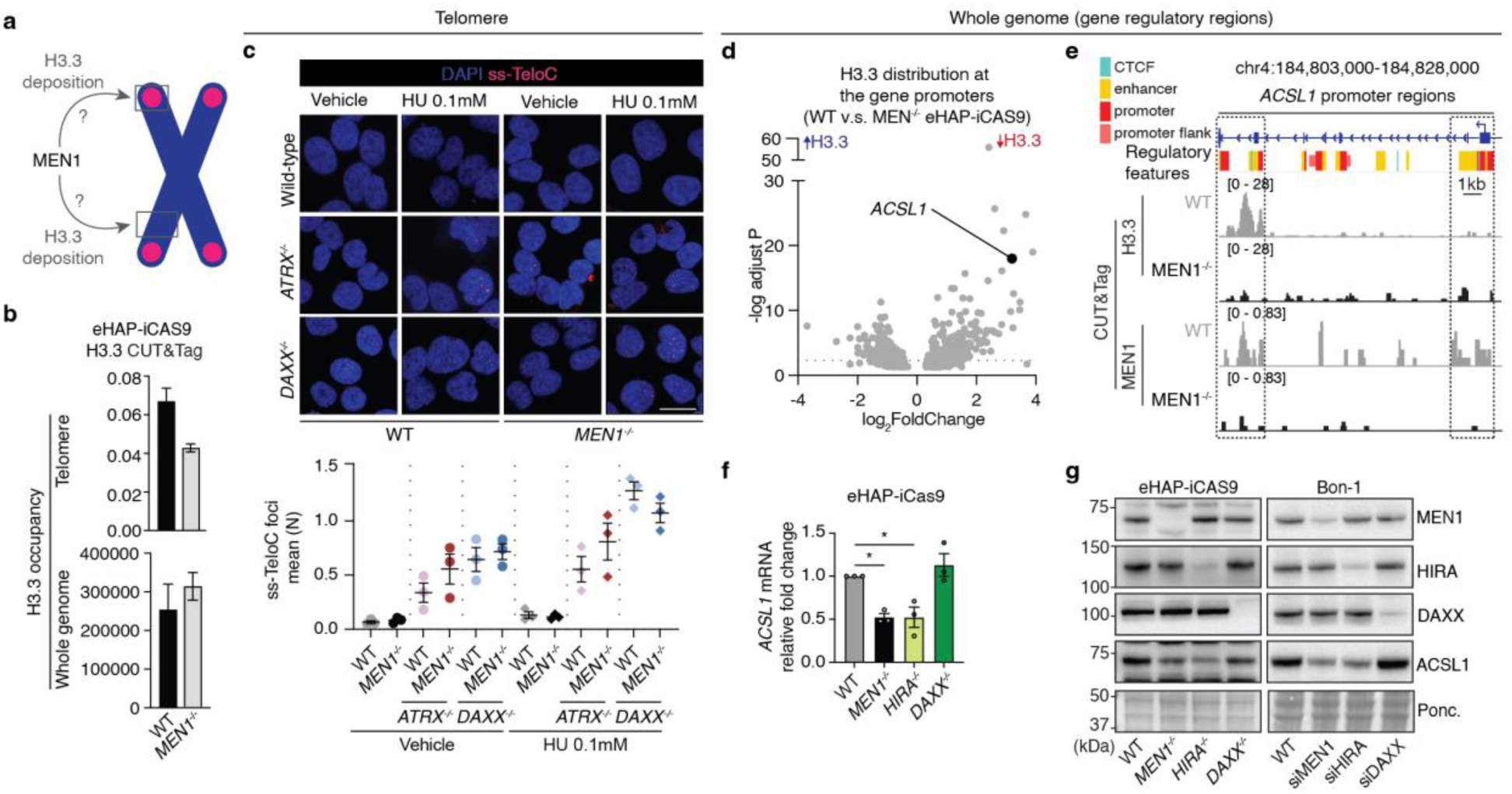
MEN1 regulates ACSL1 expression through H3.3 deposition. **a**, Schematic illustrating the dual role of H3.3 deposition. ATRX/DAXX-mediated H3.3 incorporation stabilises telomeres, while HIRA-mediated H3.3 deposition at gene-regulatory elements supports transcription. MEN1 is depicted as a potential regulator of H3.3 dynamics at both telomeres and gene loci (question marks indicate the relationships under investigation). **b**, Quantification of H3.3 CUT&Tag occupancy across the whole genome and specifically at telomeric repeat sequences in WT and *MEN1*^-/-^ eHAP-iCas9 cells. Mean ± SEM; n =3 independent samples. **c**, Representative immunofluorescent image of native FISH ssTeloC in eHAP cells treated with 0.1mM HU for 24 hrs (top). Scale bar: 20 µm. Mean ssTeloC foci number per nucleus (bottom). Mean ± SEM; n =3 independent samples. **d**, Scatter plot of differential H3.3 CUT&Tag occupancy at promoters genome-wide, comparing WT and *MEN1*^-/-^ eHAP-iCas9 cells. The x axis shows the log2 fold change in H3.3 signal (WT relative to *MEN1*^-/-^; > 1 indicates reduction in *MEN1*^-/-^ cells); the y axis shows the -log10 adjusted P value (DESeq2). Each dot represents one gene; *ACSL1* is highlighted as one of the genes with the most significantly reduced H3.3 occupancy at its promoter in *MEN1*^-/-^ cells. **e**, Genome browser view of the H3.3 and MEN1 CUT&Tag tracks at the ACSL1 promoter locus. Ensembl regulatory features (v115) are shown above to indicate locations of CTCF, enhancer, promoter, and promoter flank^48^. Scale bar = 1 kb. **f**, *ACSL1* mRNA levels measured by quantitative RT-PCR in WT, *MEN1*^-/-^, *HIRA*^-/-^ and *DAXX*^-/-^eHAP-iCas9 cells. Values are normalised to a housekeeping ACTB gene and expressed as relative fold change to WT. Mean ± SEM; n =3 independent experiments. **g**, Immunoblot showing ACSL1 protein levels in WT, *MEN1*^-/-^, *HIRA*^-/-^ or *DAXX*^-/-^ eHAP-iCas9 cells (left) and BON-1 cells with siRNA knockdown of MEN1, HIRA, or DAXX (right panel) and Ponceau S as loading control. ACSL1 protein expression is reduced in MEN1 and HIRA deficient cells, but not in DAXX deficient cells, implicating the HIRA chaperone in MEN1-dependent ACSL1 regulation.

As the same H3.3 pathway also operates at gene-regulatory elements, we next examined whether MEN1 loss alters H3.3 occupancy at ferroptosis-related genes. Indeed, genome-wide analysis identified ACSL1 as a locus with significantly decreased H3.3 promoter occupancy in MEN1^-/-^cells (Fig. 3d). CUT&Tag sequencing using a MEN1 antibody showed that MEN1 binds to the *ACSL1* gene regulatory regions where it colocalises with H3.3 (Fig. 3e and Extended Data Fig. 3a,3c). Analysis of gene regulatory elements revealed that the *ACSL1* locus contains multiple promoters and enhancers (Fig. 3e), among which MEN1 binds to two key regulatory regions (chr4:184824216-184827293 and chr4:184805543-184805602; Fig. 3e and Extended Data Fig. 3c). The overlap with active chromatin marks from public ChIP-seq datasets showed that the MEN1 binding to chr4:184824216-184827293 region coincides with an enrichment of H3K4me3 and H3K27ac, consistent with an active *ACSL1* promoter (Extended Data Fig. 3c). Notably, the binding of H3.3 and MEN1 to the chr4:184805543-184805602 region overlaps with H3K27ac and H3K4me1, consistent with an active enhancer (Extended Data Fig. 3c)^19,20^. Published chromatin conformation capture (Hi-C) datasets indicate that these regulatory regions also fall within a defined topologically associating domain, consistent with the notion that they are spatially proximal in three-dimensional chromatin architecture (Extended Data Fig. 3c). Additionally, we confirmed that the primary H3.3 chaperone: the HIRA complex, occupies the same regulatory domain (Extended Data Fig. 3c)^18^.

Prompted by these observations, we explored whether MEN1 and HIRA contribute to *ACSL1* expression. Indeed, *ACSL1* mRNA and protein levels were reduced in *MEN1*^-/-^ and *HIRA*^-/-^ cells, but not in *DAXX*^-/-^ cells (Fig. 3f, 3g). Similarly, siRNA knockdown of *MEN1* or *HIRA* (but not *DAXX*) reduced ACSL1 expression in PNET BON-1 cells, (Fig. 3g). Co-immunoprecipitation experiments revealed a weak but reproducible interaction between MEN1 and HIRA (Extended Data Fig. 3d). Together, these data suggest that MEN1 and HIRA contribute to *ACSL1* expression consistent with H3.3 regulation of gene expression.

### ACSL1 downregulation phenocopies MEN1-deficient cells in ferroptosis resistance

To determine whether the transcriptional consequences of MEN1 deficiency identified in isogenic cell models are detectable in patient tumours, we examined a cohort of genotyped PNETs stratified by MEN1 mutation status^28^. Expression of 34 individual genes within the KEGG Ferroptosis pathway (hsa04216) showed no consistent genotype-associated pattern (Extended Data Fig. 4a). The absence of broader single-gene associations may reflect the contribution of stromal cells and additional, unannotated genetic alterations present in *MEN1*-wild-type tumours. We therefore analysed gene expression at the level of three pre-specified functional signature groupings (iron metabolism, lipid metabolism/peroxidation and cystine/glutathione anti-oxidant metabolism, Fig. 4a and Extended Data Fig. 4a). *MEN1*-mutant tumours showed elevated cystine/glutathione antioxidant and lipid metabolism/peroxidation signature scores relative to *MEN1*-WT tumours, whereas iron metabolism was unchanged (Fig. 4a). These findings indicate that MEN1 mutation status is associated with a coordinated shift in ferroptosis-relevant gene-expression programmes in patient tumours.

**Fig. 4.**
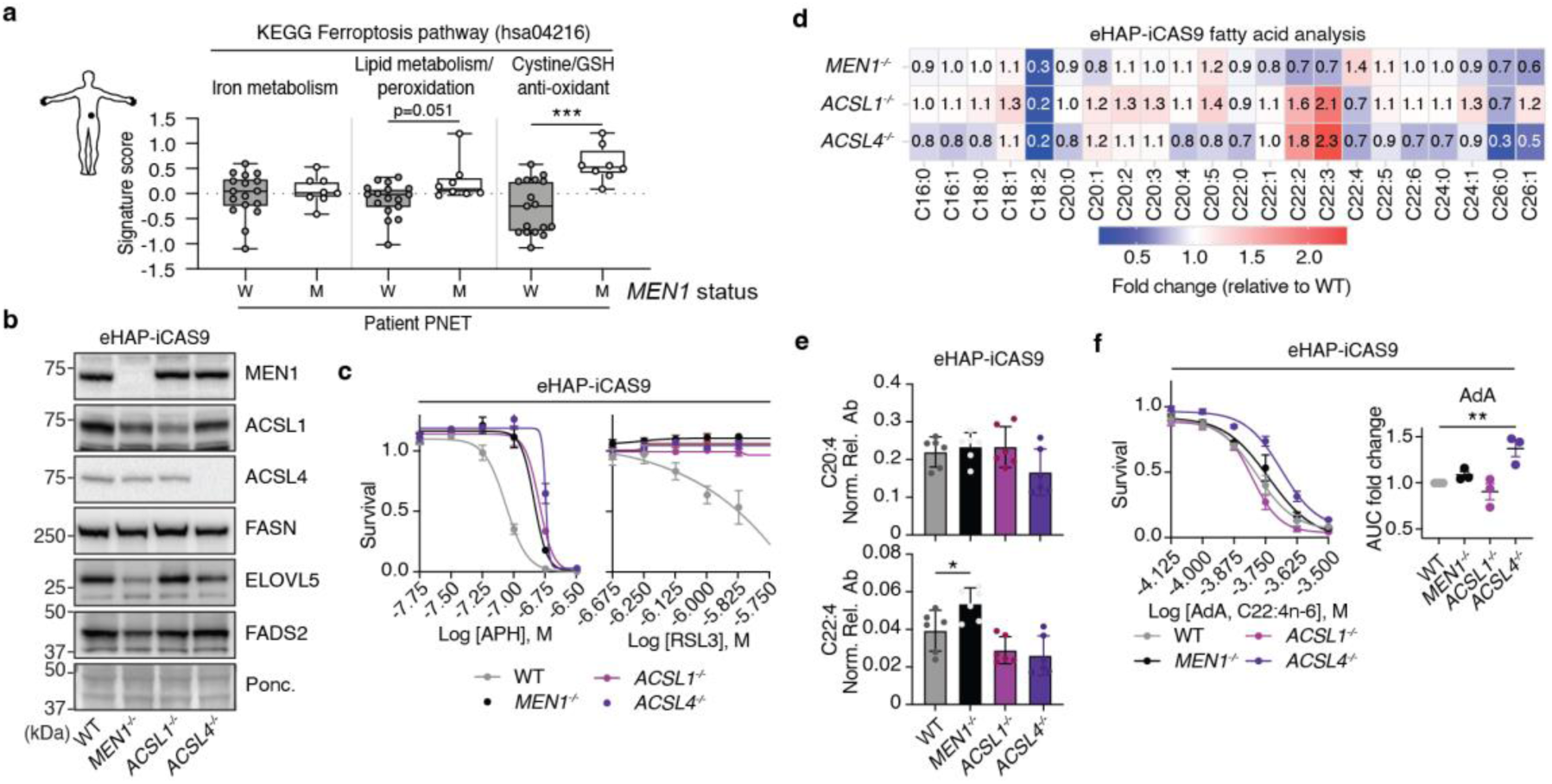
ACSL1 downregulation phenocopies MEN1-deficient ferroptosis resistance. **a,** Ferroptosis pathway signature scores differ between MEN1-wild-type (W) and MEN1-mutant (M) PNETs. A per-sample signature score was calculated as the mean expression z-score across all genes. MEN1-mutant tumours (M, n=8) showed elevated lipid metabolism/peroxidation and cystine/glutathione antioxidant signature scores relative to MEN1-wild-type tumours (WT, n=17), while no significant difference was observed for the iron metabolism. (***p<0.001; two-tailed Welch’s t-test). **b**, Immunoblot of WT, *MEN1*^-/-^, *ACSL1*^-/-^, and *ACSL4*^-/-^ eHAP-iCas9 cells probed for proteins involved in fatty acid metabolism and ferroptosis (Extended Data Fig. 4b) and Ponceau S as loading control. ELOVL5 and FADS2 are selectively reduced in *MEN1*^-/-^ cells, implicating the role of MEN1 in fatty acid regulation. **c**, Dose-response cell viability assays in eHAP-iCas9 cells of the indicated genotypes treated with APH (left) or RSL3 (right). *MEN1*^-/-^, *ACSL1*^-/-^, and *ACSL4*^-/-^ cells all show marked resistance to APH, RSL3 and erastin compared with WT. Mean ± SEM; n =3 independent experiments. **d**, Heatmap of free fatty acid relative abundances from lipidomic profiling of WT *MEN1*^-/-^, *ACSL1*^-/-^, and *ACSL4*^-/-^ eHAP-iCas9 cells, expressed as fold change relative to WT. The data represents 20 fatty acid species spanning saturated (C16:0, C18:0, C20:0, C22:0, C24:0), monounsaturated (C16:1, C18:1, C20:1, C22:1, C24:1, C26:1), and polyunsaturated (C18:2, C20:2-C20:5, C22:2-C22:6, C26:0) classes. **e**, Dot plots of normalised relative abundance for C20:4 (arachidonic acid, top) and C22:4 (adrenic acid, bottom) from lipidomic profiling of WT *MEN1*^-/-^, *ACSL1*^-/-^, and *ACSL4*^-/-^ eHAP-iCas9 cells (**d**). C20:4 is unchanged across all genotypes, indicating intact upstream PUFA biosynthesis. C22:4 is significantly elevated in *MEN1*^-/-^ cells relative to WT. (*p < 0.05; one-way ANOVA with Tukey’s post hoc test). Mean ± SEM; n =5 independent replicates.**f**, Dose-response cell viability curves for exogenous BSA-conjugated adrenic acid (AdA, C22:4n-6) across WT *MEN1*^-/-^, *ACSL1*^-/-^, and *ACSL4*^-/-^ eHAP-iCas9 cells. *MEN1*^-/-^ and *ACSL1*^-/-^ cells are not differentially affected by exogenous AdA. AUC quantification of cell viability dose-response assays in the indicated genotypes shows greater resistance to exogenous AdA in *ACSL4*^-/-^ cells compared to WT (right). Mean ± SEM; n =3 independent experiments.

### ACSL1 loss phenocopies MEN1 deficiency independently of PUFA substrate availability

ACSL1 has an established role in activating polyunsaturated fatty acids (PUFAs), which serve as substrates for phospholipid peroxidation and ferroptosis^5^. Hence, we tested whether the epigenetic downregulation of *ACSL1* could directly contribute to the observed ferroptosis resistance (Figs. 1 and 2). To this end, we generated *ACSL1*^-/-^ cells and included *ACSL4*^-/-^ cells as a control, given the established role of ACSL4 in activating PUFAs (Fig. 4b)^9^. Importantly, we found that *ACSL1*^-/-^cells phenocopied the resistance of *MEN1*^-/-^ cells in dose-response survival assays with APH or RSL3 (Fig. 4c). *ACSL4*^-/-^ cells also exhibited resistance (Fig. 4c), as previously reported^9^.

ACSL1 and ACSL4 activate long-chain fatty acids to acyl-CoA thioesters for incorporation into membrane phospholipids. ACSL4 preferentially converts PUFAs such as arachidonic acid (C20:4) and adrenic acid (C22:4) into peroxidation-prone phosphatidylethanolamine species, whereas ACSL1 exhibits a broader substrate specificity that influences both lipid remodelling and downstream partitioning (Ext Data Fig. 4b)^5^. To determine whether the ferroptosis resistance arises from changes in the lipid landscape, we performed targeted fatty acid metabolomic analysis in WT, *MEN1*^-/-^, *ACSL1*^-/-^ and *ACSL4*^-/-^ eHAP-iCas9 cells (Fig. 4d). Most fatty acids implicated in ferroptosis, including C20:4, remained unchanged, indicating intact upstream PUFA biosynthesis (Fig. 4d, 4e). However, pro-ferroptotic adrenic acid (C22:4) accumulated selectively in *MEN1*^-/-^cells, but not in *ACSL1*^-/-^ or *ACSL4*^-/-^ cells (Fig. 4e). We separately examined ELOVL5 and FADS2, which catalyse the final steps of ω-6 PUFA elongation and desaturation upstream of C22:4^29,30^; both enzymes were reduced in MEN1-deficient cells (Fig. 4b and Extended Data Fig. 4c). Because this analysis reports the free fatty acid pool, and because reduced ELOVL5 and FADS2 would be expected to lower rather than raise C22:4 production, the accumulation is most consistent with reduced utilisation of C22:4, its activation to acyl-CoA and incorporation into membrane phospholipids, rather than with increased synthesis. That *ACSL1*⁻/⁻ cells showed no such accumulation (Fig. 4e) indicates this is not simply a consequence of reduced ACSL1, and points to a broader influence of MEN1 on lipid metabolism.

To further characterise the lipidomic changes associated with MEN1 and ACSL1 deficiency, we examined additional long-chain and very-long-chain fatty acids. Most C22 species (C22:0, C22:1, C22:5 and C22:6) showed no significant differences across genotypes (Extended Data Fig. 4d). In contrast, C22:2 and C22:3 were elevated in ACSL1^-/-^ and ACSL4^-/-^ cells relative to WT and MEN1^-/-^ cells. Analysis of C26 fatty acids revealed a selective reduction of C26:0 and C26:1 in ACSL4^-/-^ cells, with intermediate levels in MEN1^-/-^ and ACSL1^-/-^ cells (Extended Data Fig. 4e). These data indicate that loss of MEN1 or ACSL1 produces a restricted set of changes in the free fatty acid pool, with the most prominent alteration being the accumulation of free adrenic acid (C22:4) specifically in MEN1^-/-^ cells.

Adrenic acid is a precursor of peroxidation-prone phosphatidylethanolamine species (PE-C22:4) that drive ferroptosis^31^. If MEN1 loss impairs C22:4 activation, *MEN1*⁻/⁻ cells should therefore resist an adrenic acid challenge. We treated WT, *MEN1*^-/-^ and *ACSL1*^-/-^ cells with BSA-conjugated adrenic acid, which reduced viability in WT cells, confirming effective uptake and the expected toxicity (Fig. 4f). Contrary to this prediction, *MEN1*^-/-^ and *ACSL1*^-/-^ cells were not protected (Fig. 4f). *ACSL4*^-/-^ followed the predicted pattern of resistance, consistent with the role of ACSL4 in driving lipid peroxidation through activation or acyl-CoA formation/acylation. These results indicate that ferroptosis resistance conferred by MEN1 or ACSL1 deficiency is not primarily dependent on changes in lipid metabolism or substrate availability.

### Selenite drives GPX4-dependent ferroptosis resistance in MEN1-deficient cells

Prompted by the elevated Cystine/GSH antioxidant signature score observed in MEN1-mutant PanNETs (Fig. 4a), we next examined whether MEN1 deficiency alters key components of the glutathione-dependent ferroptosis defence pathway. Previous studies have established that antioxidant pathways suppressing lipid peroxidation, particularly GPX4 and the parallel ferroptosis regulators GPX1 and FSP1, are critical determinants of ferroptosis sensitivity^10,25^. In eHAP-iCas9 cells, MEN1 or ACSL1 deficiency was associated with an increase in GPX4 protein levels, whereas GPX1 and FSP1 remained unchanged (Extended Data Fig. 5a). Surprisingly, this pattern was not recapitulated in BON-1 cells, where MEN1 or ACSL1 depletion did not alter GPX4, GPX1 or FSP1 expression. Instead, we noted that BON-1 cells displayed markedly lower basal levels of the selenoproteins GPX4 and GPX1 compared with eHAP-iCas9 cells (Extended Data Fig. 5a).

Because environmental factors are increasingly recognised as important modulators of ferroptosis sensitivity^12,13,32^, we considered whether differences in cell culture conditions might account for these distinct changes in eHAP and BON-1 cells. A systematic comparison of the culture media used for each model identified sodium selenite as a notable component present in IMDM (used for eHAP cells) but absent from DMEM (used for BON-1 cells) (Fig. 5a). Selenite is a key substrate for selenoprotein biosynthesis and supports expression of ferroptosis suppressors including GPX4 and GPX1^13,32^. Even trace amounts of sodium selenite in serum or standard culture media can alter GPX4 abundance and ferroptosis sensitivity, underscoring the importance of micronutrient composition in experimental systems^14^. Consistent with this, supplementation of selenite-free DMEM with sodium selenite at the concentration present in IMDM increased GPX4 and GPX1 expression in both eHAP-iCas9 and BON-1 cells (Fig. 5c).

**Fig. 5.**
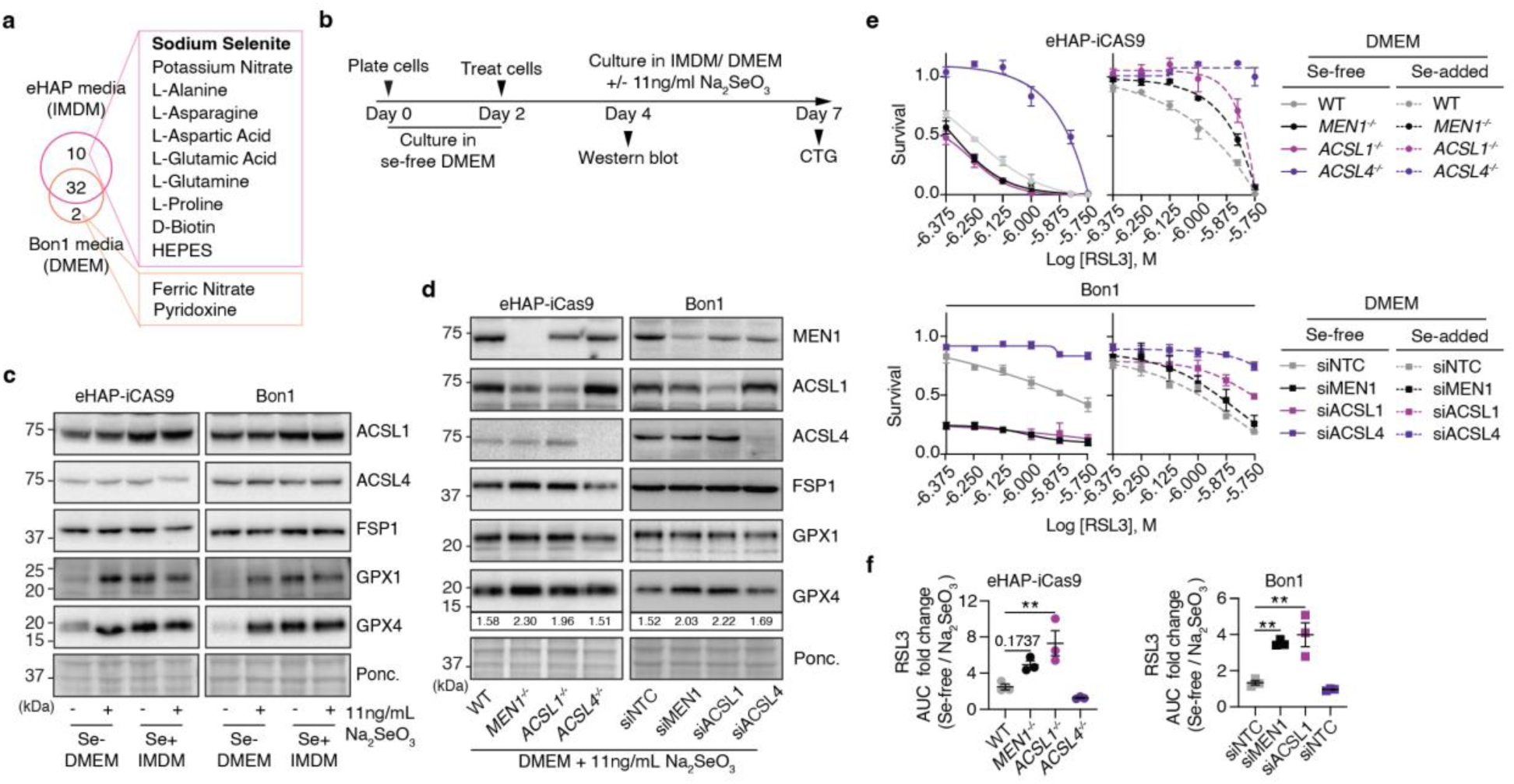
Selenite unmasks a ferroptosis-sensitised state in MEN1- and ACSL1-deficient cells. **a**, Venn diagram comparing the components of IMDM (eHAP-iCas9 culture medium) and DMEM (BON-1 culture medium). **b**, Experimental timeline for selenite treatment assays. Cells are plated on Day 0 in selenite-free DMEM, treated with the indicated compounds, and subsequently cultured in IMDM or DMEM ± 11 ng/mL Na₂SeO₃. Endpoint readouts are immunoblot (protein analysis) and CellTiter-Glo (CTG) luminescent cell viability assay. **c**, Immunoblot of eHAP-iCas9 (left) and BON-1 (right) cells cultured in the indicated selenite conditions (Se-free DMEM, Se-free IMDM, IMDM, or DMEM supplemented with 11 ng/mL Na₂SeO₃), probed for proteins involved in ferroptosis. GPX1 and GPX4 protein levels are induced by selenite in both eHAP-iCas9 and BON-1 cells. ACSL1, ACSL4, and FSP1 levels are unchanged across selenite conditions, confirming that selenite selectively induces selenoproteins without broadly altering ferroptosis pathway components. **d**, Immunoblot of eHAP-iCas9 cells of the indicated genotypes (WT *MEN1*^-/-^, *ACSL1*^-/-^, *ACSL4*^-/-^) in DMEM supplemented with 11 ng/mL Na₂SeO₃, probed for proteins involved in ferroptosis and Ponceau S (loading control). Band intensity quantification (normalized to Ponceau S and WT condition) are shown below respective protein bands. GPX4 is induced by selenite across all genotypes, with proportionally greater induction in *MEN1*^-/-^ and *ACSL1*^-/-^ cells. **e**, RSL3 dose-response cell viability assays in eHAP-iCas9 cells (top) and BON-1 cells under siRNA knockdown (bottom), cultured in selenite-free DMEM (left) or selenite-supplemented DMEM (11 ng/mL Na₂SeO₃, right). In selenite-free conditions, MEN1 and ACSL1 deficient cells are hypersensitive to RSL3 relative to WT or siNTC controls, revealing an underlying ferroptosis-sensitised state. Mean ± SEM; n =3 independent experiments. **f**, AUC fold change (selenite-free / sodium selenite-supplemented) for eHAP-iCas9 (top) and BON-1 (bottom) cells across the indicated conditions. Selenite supplementation fully restores and exceeds WT -level resistance in MEN1 and ACSL1 deficient cells but not in ACSL4 deficient cells. Mean ± SEM; n =3 independent experiments.

We proceeded to explore the impact of selenite and found that its addition significantly affects the response to ferroptosis induction with RSL3 (GPX4 inhibitor). In selenite-free conditions, *MEN1*^-/-^ and *ACSL1*^-/-^ eHAP-iCas9 cells became hypersensitive to RSL3 relative to WT cells, revealing an underlying ferroptosis-sensitised state (Fig. 5e). The same selenite-dependent pattern was recapitulated in BON-1 cells upon siRNA knockdown of MEN1 or ACSL1, but not ACSL4 (Fig. 5e). In the presence of selenite, both MEN1- and ACSL1-deficient eHAP-iCas9 or BON-1 cells became highly resistant to RSL3 as shown by the significantly increased fold change in the area-under-curve (Fig. 5f). This is consistent with specific selenite-induced GPX4 expression in MEN1-and ACSL1-deficient cells (Fig. 5d). Given the pronounced differential sensitivity to RSL3 between selenite-supplemented and selenite-free conditions, we tested whether the protective effect also involves the parallel ferroptosis suppressor FSP1^10^. Given the role of FSP1 as a secondary antioxidant involved in selenite induced ferroptosis resistance^33^, we found that it did not affect cell viability under any condition (Extended Data Fig. 5b,5c). Consistently, no change was observed in FSP1 protein levels post-selenite treatment (Fig. 5c,5d), confirming that the observed ferroptosis protection is FSP1-independent. These data suggest that the replication stress and ferroptosis resistance in MEN1- and ACSL1-deficient cells is highly dependent on GPX4 expression level when selenite is present (Fig. 5d).

### Reduced SLC7A11 and glutathione depletion create an intrinsic ferroptosis vulnerability in MEN1- and ACSL1-deficient cells

Having demonstrated that selenite availability determines the outcome of MEN1 or ACSL1 deficiency, we next examined how these perturbations intersect with the canonical SLC7A11-glutathione-GPX4 axis^34^. Notably, we found that SLC7A11 was reduced in both MEN1- and ACSL1-deficient cells (Fig. 6a), consistent with the reported role of ACSL1 in regulation of SLC7A11 protein^8^. Moreover, silencing SLC7A11 failed to alter the increase in GPX4 observed in MEN1- and ACSL1-deficient cells, suggesting that GPX4 regulation operates through a pathway distinct from the canonical SLC7A11 axis (Fig. 6a, 6b and Extended Data Fig. 6a,b).

**Fig. 6.**
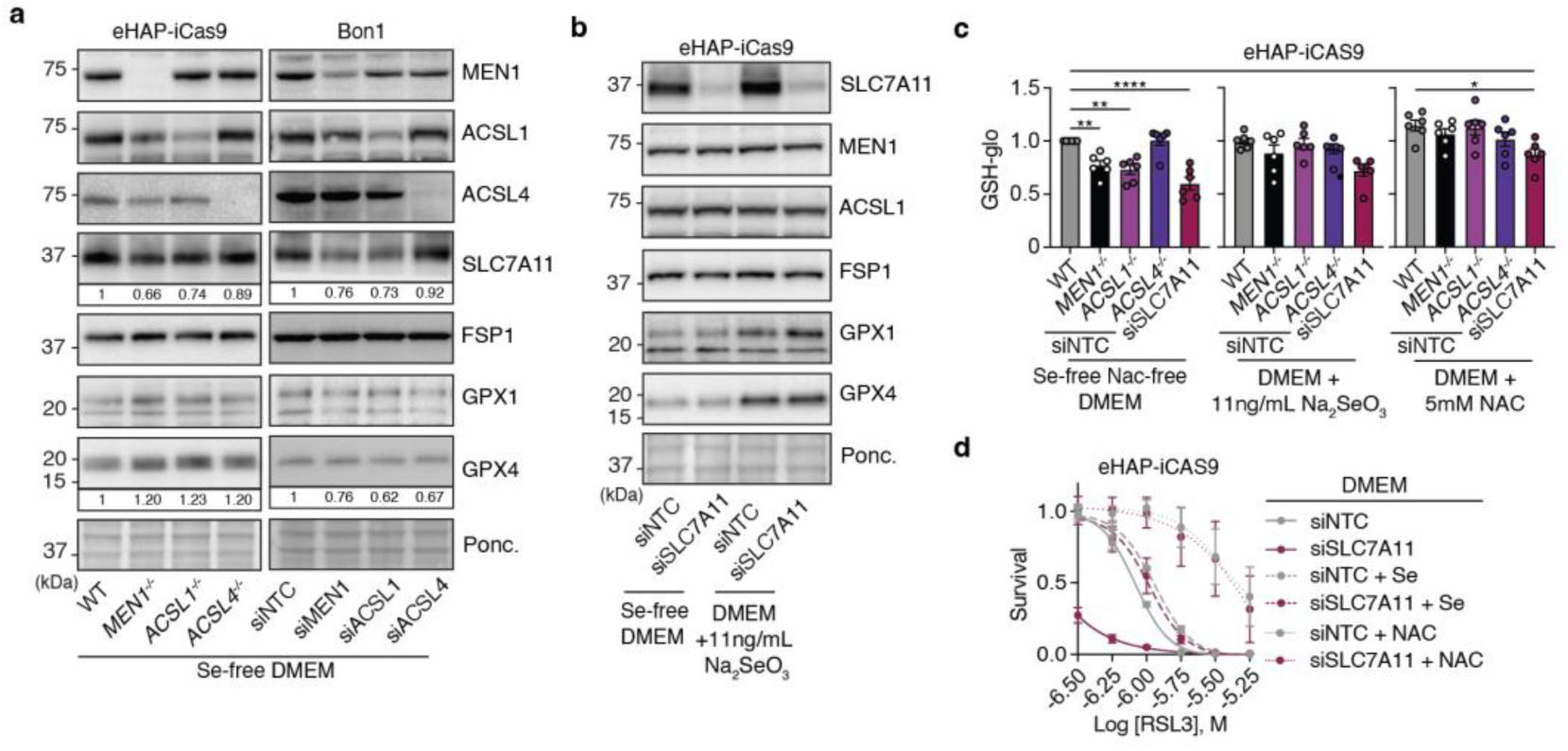
Selenite drives SLC7A11-independent GPX4 upregulation that overrides the antioxidant deficit. **a**, Immunoblot of eHAP-iCas9 cells of the indicated genotypes (WT *MEN1*^-/-^, *ACSL1*^-/-^, *ACSL4*^-/-^) in selenite-free DMEM, probed for proteins involved in ferroptosis and Ponceau S (loading control). Band intensity quantification (normalized to Ponceau S and WT condition) are shown below respective protein bands. SLC7A11 protein levels are reduced in *MEN1*^-/-^ and *ACSL1*^-/-^ cells; quantification values relative to WT are shown below the SLC7A11 blot. **b**, Immunoblot of eHAP-iCas9 cells under siRNA knockdown of SLC7A11 (siSLC7A11) or non-targeting control (siNTC) in selenite-free DMEM or DMEM supplemented with 11 ng/mL Na₂SeO₃, probed for proteins involved in ferroptosis and Ponceau S (loading control). GPX4 is induced similarly by selenite in both siNTC and iSLC7A11 cells, demonstrating that selenite-driven GPX4 upregulation is independent of SLC7A11. **c**, GSH-Glo luminescent assay quantifying intracellular glutathione (GSH) in eHAP-iCas9 cells of the indicated genotypes across three conditions: selenite-free NAC-free DMEM (left), DMEM + 11 ng/mL Na₂SeO₃ (center), and DMEM + 5 mM N-acetylcysteine (NAC, right). Values are normalised to WT controls. GSH is significantly reduced in *MEN1*^-/-^, *ACSL1*^-/-^ and siSLC7A11 cells in selenite-free conditions. Selenite and NAC supplementation restore GSH levels. (*p < 0.05, **p < 0.01, ***p < 0.001, ****p < 0.0001; one-way ANOVA with Tukey’s post hoc test). Mean ± SEM; n =3 independent experiments. **d**, RSL3 dose-response cell viability assays in eHAP-iCas9 cells cultured in selenite-free DMEM under the indicated siRNA and supplementation with 11 ng/mL Na₂SeO₃ or 5 mM N-acetylcysteine (NAC). siSLC7A11 cells acquire RSL3 resistance specifically upon selenite supplementation, phenocopying *MEN1*^-/-^ and *ACSL1*^-/-^ cells. NAC supplementation rescues RSL3 sensitivity. Mean ± SEM; n =3 independent experiments.

Nevertheless, we were intrigued by the high sensitivity of BON-1 cells to RSL3 treatment in MEN1- and ACSL1-deficient cells in selenite-free media, contrary to resistance observed in the Se-present media (Fig. 5e). Since we observed that SLC7A11 expression was downregulated in these conditions, we also checked the GSH levels considering the SLC7A11’s role in cystine import^35^. We confirmed that intracellular glutathione was significantly reduced in MEN1, ACSL1 and SLC7A11 deficient cells, and this deficit was rescued by N-acetylcysteine (NAC) supplementation (Fig. 6c and Extended Data Fig. 6c). We also found that selenite restores GSH levels, which correlates with GPX4 induction (Fig. 5c). This may reflect reduced oxidative consumption or broader selenoprotein-dependent redox adaptation (Fig. 6c and Extended Data Fig. _6c)13,36._

Consistent with the reduced SLC7A11 expression in MEN1- and ACSL1-deficient cells, SLC7A11 knockdown alone phenocopied the selenite-dependent ferroptosis response observed following MEN1 or ACSL1 depletion (Fig. 6d and Extended Data Fig. 6d). However, selenite supplementation restored RSL3 resistance in SLC7A11-deficient cells only to WT levels and failed to reproduce the enhanced resistance observed in MEN1- and ACSL1-deficient cells (Fig. 6d). This finding aligns with the preferential GPX4 induction seen specifically in MEN1- and ACSL1-deficient conditions.

Given the established role of SLC7A11 in cystine import, we next tested whether replenishing intracellular cysteine could rescue ferroptosis sensitivity. Indeed, NAC supplementation restored viability in SLC7A11-deficient cells, as well as in MEN1- and ACSL1-deficient cells where SLC7A11 expression is reduced (Fig. 6d and Extended Data Fig. 6d,e). This rescue is consistent with NAC bypassing defective cystine uptake and replenishing intracellular antioxidant capacity. Because NAC can additionally function as a ROS scavenger, its protective effect may also contribute to suppression of RSL3-induced oxidative stress, consistent with our earlier observations linking MEN1 to ROS-associated ferroptosis responses (Fig. 1e).

Together, these findings support a model in which MEN1 and ACSL1 deficiency reduce SLC7A11 expression, lowering glutathione-dependent antioxidant capacity and establishing a cystine limited, ferroptosis-sensitised state. Selenite counteracts this vulnerability through GPX4 upregulation via a mechanism operating independently of the canonical SLC7A11-glutathione pathway, restoring antioxidant protection and promoting ferroptosis resistance. MEN1 and ACSL1 deficiency therefore establishes a context-dependent binary ferroptosis response: an intrinsic vulnerability revealed under selenite-limiting conditions, and a resistant state that emerges in the presence of selenite (Fig. 7).

**Fig. 7.**
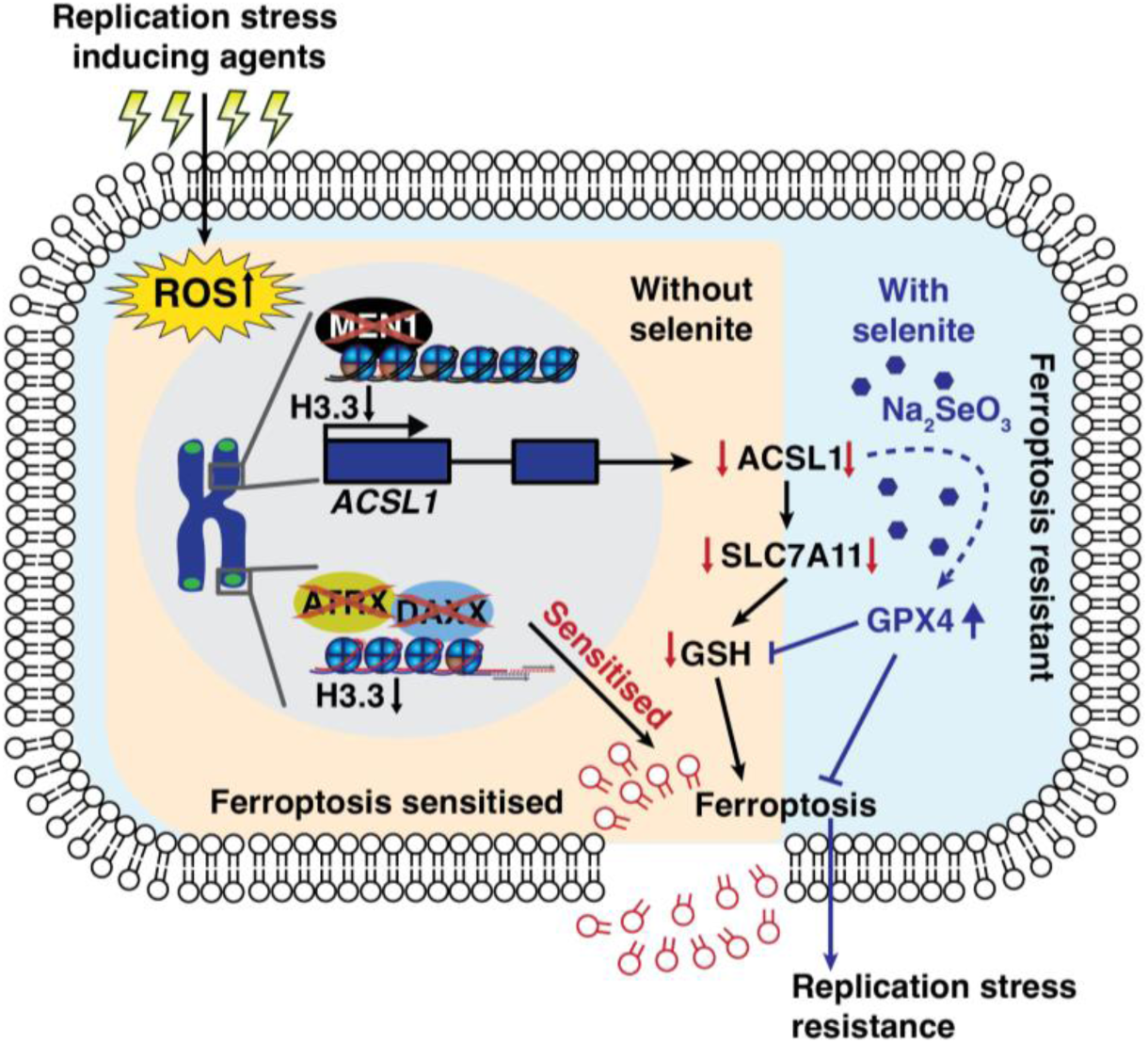
MEN1 deficiency creates a selenite-dependent switch in ferroptosis. A schematic model of the selenite-dependent switch governing ferroptosis in MEN1-deficient cells. MEN1 deficiency, acting through the HIRA-H3.3 epigenetic axis, reduces ACSL1 expression, which in turn reduces SLC7A11 and reduces intracellular GSH. In the absence of selenite, this lowered antioxidant capacity renders cells hypersensitive to GPX4 inhibition (ferroptosis-sensitised state). When environmental selenite (Na₂SeO₃) is present, it drives GPX4 upregulation through an SLC7A11-independent route (blue arrow), overcoming the GSH deficit and conferring ferroptosis and replication stress resistance.

## Discussion

This study establishes MEN1 as a key gatekeeper connecting replication stress to ferroptotic cell death. Rather than altering primary DNA-damage signalling or fork protection, MEN1 deficiency acts downstream of ROS generation to selectively impair lipid peroxidation^23^. This decouples replicative damage from death execution, conferring resistance across distinct genetic contexts, including ATRX- and DAXX-deficient backgrounds. More broadly, these findings expand the conceptual framework of replication-stress tolerance beyond checkpoint signalling and lesion repair, establishing the cell’s metabolic death machinery as a non-canonical layer of genome-stability control.

Mechanistically, we show that MEN1 functions through a HIRA-associated pathway at active regulatory elements to sustain ACSL1 expression, with direct functional consequences for lipid metabolism and ferroptosis. ACSL1 loss phenocopied MEN1 deficiency during both aphidicolin treatment and direct GPX4 inhibition, linking this transcriptional program to replication stress tolerance and GPX4-dependent ferroptosis resistance. To determine whether this protection stems from altered lipid composition, we profiled fatty acid dynamics. However, lipid substrate depletion did not drive resistance to GPX4 inhibition. Most ferroptosis-relevant fatty acids were unchanged, and adrenic acid accumulation in MEN1-deficient cells did not protect against exogenous adrenic acid challenge. MEN1 loss is additionally associated with reduced levels of other key enzymes in lipid metabolism, including ELOVL5 and FADS2, indicating a broader metabolic scope. Nevertheless, the most robust and consistent effect across both MEN1 and ACSL1 deficiency was the depletion of SLC7A11 and glutathione. This establishes that the MEN1-ACSL1 axis supports resistance to GPX4-mediated ferroptosis primarily through antioxidant capacity rather than fatty acid metabolism alone.

This vulnerability of MEN1 or ACSL1 deficiency proved conditional on selenite availability. Under selenite-limiting conditions, reduced SLC7A11 and glutathione left MEN1- and ACSL1-deficient cells hypersensitive to GPX4 inhibition. Selenite supplementation reversed this phenotype by preferentially inducing GPX4 through a route independent of SLC7A11 and FSP1, converting these same cells to a level of resistance exceeding that of wild-type controls. A single genetic lesion can therefore sensitize or protect depending entirely on a trace nutrient in the medium. This finding helps reconcile conflicting reports of ACSL1 as pro- or anti-ferroptotic across different tumour contexts and underscores that culture-medium composition is a fundamental biological variable, not merely a technical detail.

The influence of MEN1 on H3.3 is not confined to gene-regulatory elements. At the chromatin level, we establish a role for MEN1 as a regulator of histone variant H3.3 deposition. This compartmentalized control of H3.3 dynamics provides a potential mechanistic explanation for why MEN1 alterations frequently co-occur with ATRX or DAXX mutations in pancreatic neuroendocrine tumours. As cancer progression frequently relies on cooperating lesions, our data raise the possibility that MEN1 and ATRX/DAXX loss cooperate at the chromatin level. Specifically, MEN1 loss reduced telomeric H3.3 occupancy but was insufficient on its own to produce the full alternative lengthening of telomeres (ALT) phenotype observed upon ATRX or DAXX loss. Although previous whole-genome studies linked MEN1 loss to increased telomere length and ALT, these cases frequently carried co-occurring ATRX or DAXX mutations¹⁷; in contrast, isolated MEN1 loss was insufficient to establish ALT under the conditions tested here. MEN1 loss may confer a permissive chromatin state at telomeres that requires subsequent ATRX or DAXX loss to fully manifest the ALT phenotype, ultimately accounting for their frequent genetic co-occurrence in patient tumours.

Together, these findings establish MEN1 as a chromatin regulator of ferroptotic cell fate whose functional output becomes visible only when micronutrient availability is accounted for. Whether MEN1 loss sensitises cells to or protects them from ferroptosis induction therefore depends on selenite availability rather than on genotype alone. This insight carries implications beyond MEN1-mutant malignancies: any genetic regulator of ferroptosis evaluated under standard culture conditions may reflect a nutrient-dependent state rather than a fixed, immutable property of the mutation itself. Ultimately, defining how chromatin regulators intersect with micronutrient availability will be necessary to accurately map genetic dependencies in ferroptosis, and to identify which specific contexts render this pathway a viable therapeutic target in MEN1-mutant and other chromatin-altered cancers.

## Methods

### Cell culture and genome editing

#### Cell lines

The eHAP-inducible Cas9 (iCas9) was generated previously and maintained in Iscove’s modified Dulbecco’s medium (Gibco 12440053) supplemented with 10% tetracycline-free fetal bovine serum (FBS; Pan Biotech P30-3602) and penicillin-streptomycin (Gibco 15140122)^21^. The human pancreatic neuroendocrine tumour cell line BON-1 was a gift from Prof. Martyn Caplin (UCL) and maintained in Dulbecco’s modified Eagle’s medium (Gibco 41966029) supplemented with 10% tetracycline-free FBS and penicillin-streptomycin. All cell lines were grown at 37 °C in 5% CO₂ and routinely tested negative for mycoplasma. For experiments supplemented with 11 ng/mL sodium selenite (Sigma S9133) or 5 mM N-acetylcysteine (NAC; Sigma, A9165), eHAP-iCas9 is cultured in DMEM for 4 days, before treatment with sodium selenite or NAC.

#### Generation of CRISPR knockout cell lines

Full gene knockouts were generated in eHAP-iCas9 cells by reverse transfection of synthetic crRNA:tracrRNA duplexes (Integrated DNA Technologies) complexed with Lipofectamine RNAiMAX (Thermo Fisher Scientific, 13778150). Cas9 expression was induced with 1 μg ml⁻¹ doxycycline (Sigma, D9891) for 72 h post-transfection. Single-cell clones were derived by limiting dilution in 96-well plates and validated by immunoblotting. The following Edit-R crRNAs (Dharmacon/Horizon Discovery) were used: MEN1 (CM-011082-05-0002), HIRA (CM-013610-05-0002), ACSL1 (CM-011654-05-0002), and ACSL4 (CM-009364-02-0002). ATRX⁻/⁻ and DAXX⁻/⁻ lines were generated as previously described^21^. All knockout clones were validated by immunoblotting for deficiency of the target protein.

#### siRNA transfection

ON-TARGETplus SMARTPool siRNA (Horizon) were resuspended in siRNA dilution buffer (Dharmacon, B-002000-UB-100) to a concentration of 20 μM. For transient knockdown, BON-1 cells or eHAP-iCas9 were forward-transfected with ON-TARGETplus siRNA SMARTpools (Dharmacon/Horizon Discovery) targeting MEN1 (L-011082-00-0005), HIRA (L-013610-00-0005), DAXX (L-004420-00-0005), ACSL1 (L-011654-00-0005), ACSL4 (L-009364-00-0005), SLC7A11 (L-007612-01-0005), or non-targeting control siRNA (D-001810-10-05) using Lipofectamine RNAiMAX (Thermo Fisher Scientific, 13778150) according to the manufacturer’s instructions. Knockdown efficiency was confirmed by immunoblotting 72 hr post-transfection.

### Cell viability and biochemical assays

#### Colony formation viability assay

For clonogenic survival assay, 200 eHAP-iCas9 cells per well were seeded in 24-well plates in technical duplicate and cultured for 5 days in the presence of indicated drugs. Colonies were fixed, stained with 0.5% crystal violet in 20% methanol, imaged and quantified using GelCount software (Oxford Optronics). Colony numbers were normalised to untreated WT controls.

#### Treatments

For luminescence-based viability assays, eHAP-iCas9 cells were seeded at 200 cells per well (5-day assays) in opaque white 384-well plates or 5,000 cells per well (24 h endpoint assays) in opaque white 96-well plates (Greiner). BON-1 cells were seeded at 4,000 cells per well (5-day assays) in opaque white 384-well plates or 20,000 cells per well for 24 h endpoints. Compounds used included hydroxyurea (Sigma, H8627), aphidicolin (Sigma, A0781), RSL3 (Selleckchem, S8155), erastin (Selleckchem, S7242), ferrostatin-1 (Sigma, SML0583), ATR inhibitor AZD6738 (Selleckchem, S7693), Wee1 inhibitor MK-1775 (Selleckchem, S1525), A-1331852 (Selleckchem, S7801), ABT-199 (Selleckchem, S8048), ABT-263 (Selleckchem, S1001), and S63845 (Selleckchem, S8383). For multi-day assays, cell survival was assessed after 5 days. For 24-hour endpoint assays, cell viability was measured 24 hours post-treatment. In both cases, viability was quantified using the CellTiter-Glo Luminescent Cell Viability Assay (Promega, G7570) according to the manufacturer’s instructions, and luminescence was measured on a CLARIOstar microplate reader (BMG Labtech). Luminescence values were normalised to wells with vehicle control for each cell line. Area under the curve (AUC) was calculated from dose-response data and expressed as fold change relative to the WT or siNTC control.

#### BSA-fatty acid conjugation and exogenous fatty acid treatment

Fatty acid-BSA conjugates were prepared fresh before each experiment. Bovine serum albumin (Sigma, A8806) was dissolved in serum-free IMDM at 55 °C to a concentration of 10% (w/v) with gentle agitation. Adrenic acid (cis-7,10,13,16-docosatetraenoic acid; Sigma, D3659) were dissolved in 0.1 M NaOH at 70 °C to a concentration of 20 mM. These fatty acid solutions were then slowly mixed with the BSA solution at a 6:1 molar ratio (10 mM fatty acid : 1.7 mM BSA), stirred for 1 h at 37 °C, pH-adjusted to 7.4, and sterile-filtered. Matched BSA vehicle controls were prepared identically without fatty acid and added at the same concentrations. For fatty acid supplementation assays, eHAP-iCas9 cells (5,000 per well) or BON-1 cells (20,000 per well) were seeded in opaque white 96-well plates (Greiner), allowed to adhere for 24 h, and then treated with the BSA-conjugated fatty acids at the indicated concentrations for 24 h. Cell viability was assessed by CellTiter-Glo Luminescent Cell Viability Assay (Promega, G7570) as described above.

#### Quantification for ferroptosis associated biochemical markers

Reactive oxygen species (ROS) were measured using the ROS-Glo assay (Promega, G8820), lipid peroxidation by malondialdehyde (MDA) assay (Abcam, ab118970), and intracellular glutathione (GSH) using the GSH-Glo Glutathione Assay (Promega, V6911) according to the manufacturer’s instructions. Colourimetric and luminescence measurement were performed on a CLARIOstar microplate reader (BMG Labtech).

### Molecular and protein analyses

#### qRT-PCR

qRT-PCR samples were prepared using Cells-to-CT 1-Step Taqman Kit (Invitrogen) from WT MEN1^-/-^ and *HIRA*^-/-^ and *DAXX*^-/-^ eHAP-iCas9 cells. QuantStudio 5 (Invitrogen) was used for PCR with taqman probes; ACTB (Hs01060665_g1), or ACSL1 (Hs00960561_m1). Fold induction was calculated using ACTB as housekeeping gene.

#### Whole cell lysis and immunoblotting

Cells were rinsed with PBS, trypsinised, and pelleted by centrifugation at 500 × g for 5 min. Pellets were washed once with PBS and lysed in RIPA buffer (10 mM Tris-Cl pH 8.0, 1 mM EDTA, 0.5 mM EGTA, 1% Triton X-100, 0.1% sodium deoxycholate, 0.1% SDS, 140 mM NaCl) supplemented with PhosSTOP phosphatase inhibitor cocktail and cOmplete EDTA-free protease inhibitor cocktail (Roche). Lysates were clarified by centrifugation at 13,000 × g for 15 min at 4 °C. Protein concentrations were determined using the DC Protein Assay (Bio-Rad, 5000116). Proteins were resolved by SDS-PAGE on NuPAGE mini gels (Invitrogen, NP0321) and transferred to 0.2 μm nitrocellulose membranes (Amersham Protran, 10600001). Membranes were blocked in 5% skimmed milk/TBST for 1 h at room temperature, incubated with primary antibodies overnight at 4 °C, washed three times with TBST, and incubated with HRP-conjugated secondary antibodies for 1 h at room temperature. Signals were developed using Clarity or Clarity Max ECL substrate (Bio-Rad, 1705060 or 1705062) and imaged on a ChemiDoc MP system (Bio-Rad). Ponceau S staining served as loading control prior to antibody incubation.

Primary antibodies used were: anti-ATRX (Bethyl, A301-045A), anti-MEN1 (Bethyl, A300-105A), anti-DAXX (Santa Cruz, sc-7152), anti-FSP1/AIFM2 (Santa Cruz, sc-377120), anti-FASN (Abcam, ab22759), anti-xCT/SLC7A11 (Cell Signaling, 12691), anti-GPX1 (Abcam, ab22604), anti-SCD1 (Abcam, ab19862), anti-ACSL4 (Cell Signaling, 38493), anti-phospho-mTOR (Ser2448) (Abcam, ab109268), anti-ACSL1 (Thermo Fisher, 13989-1-AP), anti-ELOVL5 (Proteintech, 26599-1-AP), anti-CD44v6 (Thermo Fisher, BMS125), anti-COX2 (Cell Signaling, 12282), anti-FADS2 (Proteintech, 28034-1-AP), anti-MLKL (Merck, MABC604), anti-phospho-MLKL (Ser345) (Abcam, ab196436), anti-GPX4 (Cell Signaling, 52455), and anti-SREBP1 (Abcam, ab3259). Additional antibodies included anti-HIRA (Active Motif, 39557), anti-phospho-ATR (Ser428) (Cell Signaling, 2853), anti-γ-H2AX (Millipore, 05-636), anti-phospho-ATM (Ser1981) (Abcam, ab81292), anti-caspase-3 (Cell Signaling, 9662), anti-cleaved caspase-3 (Cell Signaling, 9664), anti-ATR (Santa Cruz, sc-1887), anti-ATM (Sigma, A1106), anti-PARP (Cell Signaling, 9542), anti-LAMP2 (Invitrogen, PA5-118026), anti-MAVS (Cell Signaling, 24930), and anti-LC3B (Abcam, ab48394).

#### Co-immunoprecipitation

For co-immunoprecipitation, eHAP-iCas9 cells were lysed in IP buffer (20 mM Tris-HCl pH 8.0, 137 mM NaCl, 1% IGEPAL, 2 mM EDTA) supplemented with phosphatase and protease inhibitor mixes (Roche) and benzonase (Sigma). Nuclei were mechanically dissociated through a 27 G syringe and incubated for 40 minutes on ice. Lysates were cleared by centrifugation at 13,000 × g for 15 minutes at 4°C. Cleared lysates were diluted in IP buffer and immunoprecipitated overnight at 4°C using protein A/G beads pre-conjugated to the indicated 2μg antibodies (anti-MEN1 or anti-HIRA) or IgG control. Beads were washed three times with dilution buffer containing 0.05% NP-40, and bound proteins were eluted in 2× NuPAGE LDS sample buffer (Invitrogen) with 1% 2-mercaptoethanol (Sigma) at 95°C.

#### Single strand-TELO C FISH assay

Fixed cells were blocked using antibody dilution buffer (ADB) consisting of 0.1% Triton X-100, 0.1% saponin, 100 μg ml⁻¹ RNase A (Thermo Fisher Scientific, EN0531), and 10% goat serum (Sigma, G9023) in PBS for 1 h at 37 °C. Cells were washed three times with PBS and dehydrated using increasing concentrations of ethanol (70%, 90%, then 96%). Once dried, 2 nM TelG TAMRA probe in hybridisation buffer (70% formamide, 1 mg ml⁻¹ Roche blocking reagent, 10 mM Tris-HCl pH 7.5) was added and incubated at room temperature for 2 h. Cells were then washed three times with PBS, counterstained with DAPI (Sigma, D9542), and stored at 4 °C before imaging on a Nikon Ti2-E inverted microscope equipped with a Nikon Plan Apo 60×/1.40 oil objective and Photometrics Prime 95B camera.

#### C-Circle assay

The C-circle assay was performed using the quick C-circle preparation (QCP) protocol^37^. Briefly, ∼2 × 10⁵ cell pellets or ∼2 mm² snap-frozen tumour samples were homogenised in 200 μL fresh QCP buffer (50 mM KCl, 10 mM Tris-HCl pH 8.5, 2 mM MgCl₂, 0.5% IGEPAL CA-630, 0.5% Tween-20) using a Precellys tissue homogeniser (Bertin Technologies). Homogenates were incubated at 56 °C with 1,400 rpm shaking for 1 h in QCP lysis buffer supplemented with 1:20 (v/v) protease (QIAGEN, 19155), followed by inactivation at 70 °C for 20 min. DNA concentration was measured by fluorimetry using the Qubit dsDNA HS Assay (Thermo Fisher Scientific, Q32854). Thirty nanograms of DNA were diluted to 10 μL in 10 mM Tris-HCl pH 8.0 and mixed with 9.25 μL rolling circle master mix (8.65 mM DTT, 2.16× phi29 buffer, 8.65 μg ml⁻¹ BSA, 0.216% Tween-20, and 2.16 mM each dATP, dCTP, dGTP, dTTP) with or without 0.75 μL phi29 DNA polymerase (Thermo Fisher Scientific, EP0094). Rolling circle amplification was performed at 30 °C for 8 h, followed by polymerase inactivation at 70 °C for 20 min. Samples were blotted onto Amersham Hybond N+ positively charged nylon membranes (GE Healthcare, RPN303B) under native conditions, UV-crosslinked, and hybridised overnight at 65 °C with 3′ DIG-labelled telomeric (5′-CTAACCCTAACCCTAACC-3′) or Alu (5′-GTAATCCCAGCACTTTGG-3′) probes prepared using the DIG oligonucleotide 3′-end labelling kit (Roche, 03353575910). Membranes were washed, incubated with anti-DIG-AP antibody (Roche, 11093274910), and developed with CDP-Star substrate (Roche, 12041677001). Chemiluminescence was acquired on a ChemiDoc MP imaging system (Bio-Rad) and quantified using Fiji/ImageJ. Membranes were stripped and rehybridised with the Alu probe as a loading control.

### Genomic and epigenomic profiling

#### CRISPR/Cas9 screening and analysis

Genome-wide CRISPR/Cas9 synthetic lethal screens were performed in eHAP-iCas9 WT or *ATRX*^-/-^ cells transduced with the lentiviral Brunello library (Addgene #73179-LV). A minimum of 100 million cells were transduced at a multiplicity of infection of 0.4 across three independent transductions, maintaining coverage of 500 cells per sgRNA. Transduced cells were selected with puromycin (0.4 μg/mL) for 48 hours, after which Cas9 was induced with 1 μg/mL doxycycline. Cells were subcultured every two days, maintaining representation of at least 40 million cells per passage. Genomic DNA was isolated using the PureLink Genomic DNA Mini Kit (Thermo Fisher Scientific), quantified by Nanodrop and Qubit, and 200 μg used for library preparation by one-step PCR amplification of integrated sgRNA sequences with Ex Taq polymerase (TaKaRa). PCR products were purified using the QIAquick Gel Extraction Kit (Qiagen), quality-controlled on Bioanalyzer (Agilent), and sequenced on HiSeq 4000 (100 bp reads, 30 million reads per sample). Raw reads were trimmed to 20 bp, mapped to the Brunello guide library using BWA (v0.5.9), and sgRNA counts generated from zero-mismatch, forward-strand alignments. The MAGeCK ‘test’ command (v0.5.7) was used for pairwise comparisons with parameters ‘--norm-method total --remove-zero both’. Dashed red line indicates the significance threshold (MAGeCK score = 1)^21^. Details of identified genes in the synthetic lethal screen in WT and *ATRX*^-/-^ are shown in Supplementary Table 1,2 respectively.

#### CUT&Tag sequencing

CUT&TAG was done as previously described ^38,39^. Briefly, eHAP cells corresponding to various genotypic conditions (WT *MEN1*^-/-^) were collected. Nuclei were extracted from each cell line using a nuclear extraction buffer (20 mM HEPES-KOH pH 7.9, 10 mM KCl, 0.5 mM spermidine, 0.1% Triton X-100, 20% glycerol, freshly added protease inhibitors). Nuclei were bound to Concanavalin A (Sigma C2272) coated MyoneT1 beads (ThermoFisher, 65601) as previously described ^40^. Bound nuclei were incubated in wash Buffer (20 mM HEPES pH 7.5; 150 mM NaCl; 0.5 mM Spermidine; 1× Protease inhibitor cocktail; 0.05% Digitonin) containing 2 mM EDTA, and a 1:50 dilution of the appropriate primary antibody containing 2 mM EDTA and a 1:50 dilution of the primary antibody targeting either H3.3 (ActiveMotif, 91191) or MEN1 (ActiveMotif, 61005) overnight at 4 °C. Bead-bound nuclei were washed and resuspended in the appropriate secondary antibody. Next, bead bound nuclei were resuspended in a Tagmentation buffer (containing pA-Tn5, 1:200, 0.04 uM) and incubated at 37 °C for exactly 1 hour. A cocktail of ProteinaseK, EDTA, and SDS were added to quench the signal and the gDNA was extracted using the Zymo extraction kit (Zymo Research. D4066). Libraries were amplified using PCR conditions and primers as previously described ^39^. Post-PCR clean-up was performed 3 times by adding 1.1× volume of Ampure XP beads (Beckman Counter). Libraries were incubated with beads for 10 min at RT, washed twice in 80% ethanol, and eluted in 10 mM Tris pH 8.0.

For bioinformatics analysis, we used a pre-established nf-core pipeline (https://nf-co.re/cutandrun/dev/docs/usage) to analyse the data. Picard was used to mark duplicates ^41^, SAMtools ^42^ to create BAM files. Since the libraries contained homemade pA-Tn5, there was residual carryover of *E.coli* DNA which was used as a spike-in. Reads were aligned to the E. coli K12-MG1655 genome and we performed a spike-in normalisation using BEDtools ^43^. SEACR ^44^ was used to call peaks with default parameters. Fragment- and peak-based quality control checks were performed using DeepTools. To plot pearsons, PCA, heatmaps, we used Deeptools (https://deeptools.readthedocs.io/en/2.2.2/index.html). Sequencing data were aligned to the human genome (GRCh38) and visualised using the Integrative Genomics Viewer (IGV, Broad Institute). For telomere analysis, *Grep* command was used to identify “TTAGGGTTAGGG” sequences in fastq, for comparison across conditions, the number of telomere sequence containing reads was normalised with the total number of reads for each condition. Details of identified genes in H3.3 occupancy comparison between WT and *MEN1*^-/-^ are shown in Supplementary Table 3. Public HIRA complex, HIRA, ASF1A, UBN1, ChIP sequencing datasets (GSE45024) were overlaid to compare HIRA complex binding at the ACSL1 promoter and enhancer regions^45^. Hi-C contact maps were obtained from GSE74072 and used to define topologically associating domains (TADs) surrounding the ACSL1 locus^46^. Regulatory features (Homo_sapiens.GRCh38.regulatory _features.v115) were downloaded from the ENCODE project consortium and used to annotate promoter, enhancer, and CTCF-bound elements together with active chromatin marks H3K4me3, H3K27ac, and H3K4me1^19^.

#### Analysis of cancer genomics data

Cancer genomics data were obtained from cBioPortal. A combined cohort of 194 samples from 178 patients with pancreatic neuroendocrine tumours was queried for somatic alterations in MEN1, ATRX, and DAXX^17,47^. Genomic profiles including mutations and copy-number alterations were analysed for co-occurrence and mutual exclusivity using default parameters. Fisher’s exact test with Benjamini-Hochberg correction was applied to calculate statistical significance (q < 0.05).

#### Analysis of patient gene expression data

Gene expression data for pancreatic neuroendocrine tumours (PNETs) genotyped for ATRX, DAXX, and MEN1 mutation status were obtained from the Gene Expression Omnibus (GEO) series GSE117851^28^, using the GEO2R analysis tool (NCBI) to extract sample-level expression values and calculate as z-score. Genotype annotations (MEN1-wild-type versus MEN1-mutant) were assigned based on mutation status reported in the original study^28^. Of the available cohort, 25 samples with unambiguous MEN1 genotype annotation were used in this analysis (MEN1-wild-type, n=17; MEN1-mutant, n=8). Functional gene grouping were defined a priori from the KEGG Ferroptosis pathway (hsa04216), by grouping the pathway-annotated genes into categories reflecting their established roles: iron metabolism, lipid metabolism and peroxidation, cystine/glutathione antioxidant metabolism, and other regulatory factors. For each tumour sample, a signature score was calculated as the mean expression z-score across all genes within that module. Signature scores were compared between MEN1-wild-type and MEN1-mutant groups using a two-tailed Welch’s t-test.

### Fatty acid analysis

#### Sample processing

For metabolite extraction, cells were harvested by gentle scraping into the culture media, transferring to a 50 mL tube and rapid metabolic quenching using a dry ice/70% ethanol slurry. Quenched cells were pelleted by centrifugation (1,100 rpm, 4 °C, 3 min), washed twice with ice-cold PBS (pH 7.3), counted, and equal cell numbers aliquoted into pre-chilled 2 mL tubes (Eppendorf, Safe-lock). After centrifugation (16,000 rpm, 4 °C, 5 min), metabolites were extracted by adding 400 μL chloroform and 200 μL Optima-grade methanol (both Fisher Scientific, 10615492 and 10767665, respectively) to the cell pellet, followed by brief vortexing, and incubation in a water bath sonicator (4 °C, 1 h, with 3 × 8 min sonication pulses). After centrifugation (16,000 rpm, 4 °C, 10 min), the supernatant was transferred to a 1.5 mL tube (Eppendorf) and dried using a rotary vacuum concentrator (Christ). The pellet was re-extracted with 450 μL methanol/water (2:1, v/v), sonicated for 8 min (water bath sonicator, 4 °C), centrifuged (16,000 rpm, 4 °C, 10 min), and the second extract combined with the (now dry) first, before drying once more. Dried extracts were resuspended in 50 μL chloroform (containing 5 nmol C17:0 internal standard), followed by addition of 150 μL methanol and 150 μL Optima-grade water (Fisher Scientific, 10505904) to achieve a final chloroform/methanol/water ratio of 1:3:3 (v/v). After vigorous vortexing and centrifugation (as above), the lower apolar phase (containing lipidic metabolites) was transferred to a glass vial insert, dried, washed twice from methanol, and derivatised to produce fatty acid methyl esters (FAMES) by addition of 25 µL chloroform/methanol (2:1, v/v) and 7 µL Trimethylphenylammonium hydroxide solution (TMAH; Merck, 79266) followed by brief vortexing and incubation (RT, 1 h).

#### Two-dimensional gas chromatography-Time of Flight mass spectrometry (GC×GC-TOFMS) analysis

Data acquisition was performed on a Pegasus BT 4D Time of Flight mass spectrometer (LECO Corp., St. Joseph, MI, USA) coupled to a 7890 gas chromatography system (Agilent Technologies) equipped with a secondary oven. Separation was achieved using a two-dimensional column set comprising an RXi-5SilMS (Restek, 30 × 0.25 mm i.d., 0.25 µm film thickness) as the first-dimension column (1D), and an RXi-17SilMS (Restek; 0.7 m × 0.25 mm i.d., 0.25 µm film thickness) as the second-dimension column (2D). The second-dimension column was connected to the TOFMS via an additional RXi-17SilMS capillary (0.1 m × 0.25 mm i.d.). Samples (1 µL) were injected in split mode (50:1) at 250 °C, with a purge time of 30 s and a purge flow of 3 mL min^-1^. Helium (99.9999% purity) was used as the carrier gas at a constant flow rate of 1.4 mL min^-1^. The primary oven temperature program was set at 70 °C (held for 1 min), then ramped at 5 °C min^-1^ to 300 °C, with a final hold of 5 min. The secondary oven was maintained at an offset of +5 °C relative to the primary oven throughout the run. Modulation was performed with a 2 s period, including 0.6 s hot pulse and 0.4 s cold pulse, with a modulator temperature offset of 15 °C. The TOFMS was operated in electron ionisation (EI) mode over a mass range of 45-650 Da. The ion source temperature was set to 250 °C, and data were acquired at 200 spectra s^-1^. Data acquisition and processing were performed using ChromaTOF software (Leco, version 5.59.51). Peak detection was performed automatically using a signal-to-noise (S/N) threshold of 100.

#### Fatty acid quantification and data analysis

Absolute quantification of derivatised fatty acids was performed using an internal standard-based approach incorporating the molar relative response factor (MRRF). The MRRF was calculated from derivatised fatty acid standard mixtures as the ratio of the mean response of each metabolite to that of the internal standard (C17:0), as below:

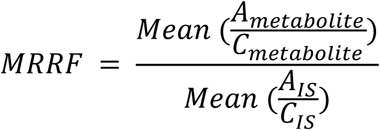

*A_metabolite_* and *A_IS_* represent the peak areas of the metabolite and internal standard (IS) in the standard mixture, respectively, and *C_metabolite_* and *C_IS_* correspond to their known amounts. The amount of each metabolite in the samples was then calculated using the ratio of the analyte peak area to that of the internal standard, multiplied by the known concentration of the metabolite in the standard mixture (5 nmol), and corrected by the MRRF:

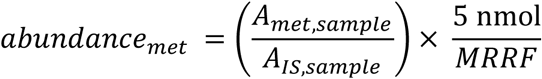

The resulting values were normalized to cell number and expressed as nmol per 10^6^ cells. Relative abundances were determined at the level of fatty acid classes defined by carbon chain length and degree of unsaturation. Fatty acid identities were confirmed by comparison to authentic standards processed in the same way and run in the same sample batch. Fatty acids for which no authentic standard was available were assigned putative annotations based on their elution position within the GC×GC chromatographic space (homologous series organization by carbon number and degree of unsaturation), characteristic mass spectral fragmentation patterns, and comparison with the NIST Mass Spectral Library. For each fatty acid class (e.g., C16, C18, C20, C22, C24, C26), peak areas corresponding to all species with the same number and degree of unsaturation, including positional isomers and putatively annotated compounds, were summed (e.g., all C20:1 species). The summed peak area of each class was normalized to the internal standard peak area within the same sample and subsequently normalized to cell number, yielding values expressed as area ratio per 10^6^ cells. Details of identified and annotated fatty acids, their peak areas, and calculated abundances are shown in Supplementary Table 4.

### Statistical analysis

The sample number (n) in each experiment indicates the number of independent biological replicates and is specified in the relevant figure legend. Statistical analysis was performed using Prism 9/10 (GraphPad) and RStudio. Unless otherwise stated, one-way ANOVA followed by Tukey’s multiple comparison test was applied. Significance thresholds: *p < 0.05, **p < 0.01, ***p < 0.001, ****p < 0.0001. Co-occurrence analysis of *MEN1*, *ATRX*, and *DAXX* mutations in PNET was performed using cBioPortal. Log2 odds ratios were calculated to quantify the tendency for co-occurrence or mutual exclusivity; Fisher’s exact test was applied, with q-values corrected by the Benjamini-Hochberg method. A q-value < 0.05 was considered statistically significant.

## Data availability

All data are available in the main text or the supplementary materials.

## Acknowledgments

We thank members of the Boulton laboratory for the discussions. We are grateful to the Francis Crick Institute’s scientific technology platforms, in particular Genomics, and Cell Services, for technical support. R.R.J.L was supported by the Horizon Europe research and innovation programme under the Marie Skłodowska-Curie grant agreement No. 101150631 and has been supported by the UKRI Horizon Europe Guarantee fund (EP/Z002648/1). Work in the S.J.B. laboratory is supported by the Francis Crick Institute (CC2057), European Research Council Advanced Investigator grants (TelMetab, ChrEndProt), a Wellcome Trust Senior Investigator Award, and Cancer Research UK RadNet City of London. The Francis Crick Institute receives its core funding from Cancer Research UK, the UK Medical Research Council, and the Wellcome Trust. We thank Paola Peinado Fernandez and Claudio Ballabio for providing feedback on the manuscript.

## Author contributions

R.R.J.L. conceived the project, designed and performed most experiments, analyzed the data. S.J.B. supervised the study and secured funding. N.P. performed CUT&Tag experiments and analysis. A.I.I. assisted with experiments and conducted C-circle assays. S.S.B. generated the genome-wide CRISPR screen dataset. E.G. and F.T.S carried out fatty acid analysis experiments and analysis with supervision from J.I.M. R.R.J.L and S.J.B wrote the manuscript with input from all co-authors.

## Materials & Correspondence

Correspondence to Ronnie Ren Jie Low and Simon J. Boulton. Unique reagents generated here are available upon request.

## Tables

Supplementary Table 1. WT untreated vs HU synthetic lethal screen

Supplementary Table 2. ATRX-KO untreated vs HU synthetic lethal screen

Supplementary Table 3. WT vs MEN1-KO H3.3 CUT&Tag promoters analysis

Supplementary Table 4. Annotated and identified fatty acid analysis

