## Supplemental figures for "MEN1 deficiency creates a selenite-dependent switch in ferroptosis"

**Extended data**

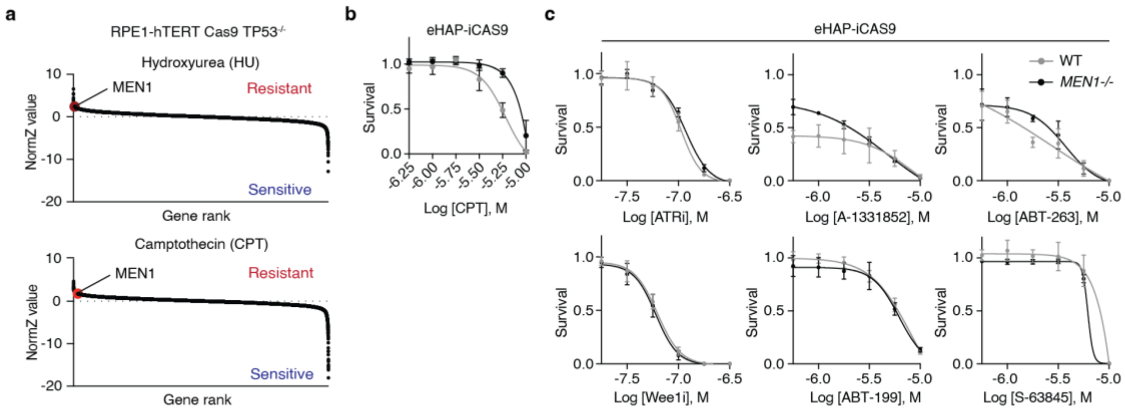


**Extended Data Fig. 1| MEN1-deficient cells show no differential sensitivity to DNA-damage-response or apoptosis inhibitors**

**a**, Scatter plots from genome-wide CRISPR-Cas9 screens performed in RPE1-hTERT Cas9 TP53$^{-/-}$ cells treated with HU (top) or camptothecin (CPT, bottom)[24]. Each dot represents one gene; MEN1 (red) ranks as a top resistance hit in both screens.

**b**, Dose-response cell viability assays in WT and *MEN1*$^{-/-}$ eHAP-iCas9 cells treated with CPT. *MEN1*$^{-/-}$ cells show resistance with CPT treatment. Mean ± SEM; n =3 independent experiments.

**c**, Dose-response cell viability assays in WT and *MEN1*$^{-/-}$ eHAP-iCas9 cells treated with ATR inhibitor AZD6738, Wee1 inhibitor MK-1775 or BH3 mimetics targeting BCL-2 (ABT-199), BCL-xL (A-1331852), MCL-1 (S63845) or BCL-2/BCL-xL/BCL-W (ABT-263) in WT and *MEN1*$^{-/-}$ eHAP-iCas9 cells. No differential response to the inhibitors to DDR or apoptosis. Mean ± SEM; n =3 independent experiments.

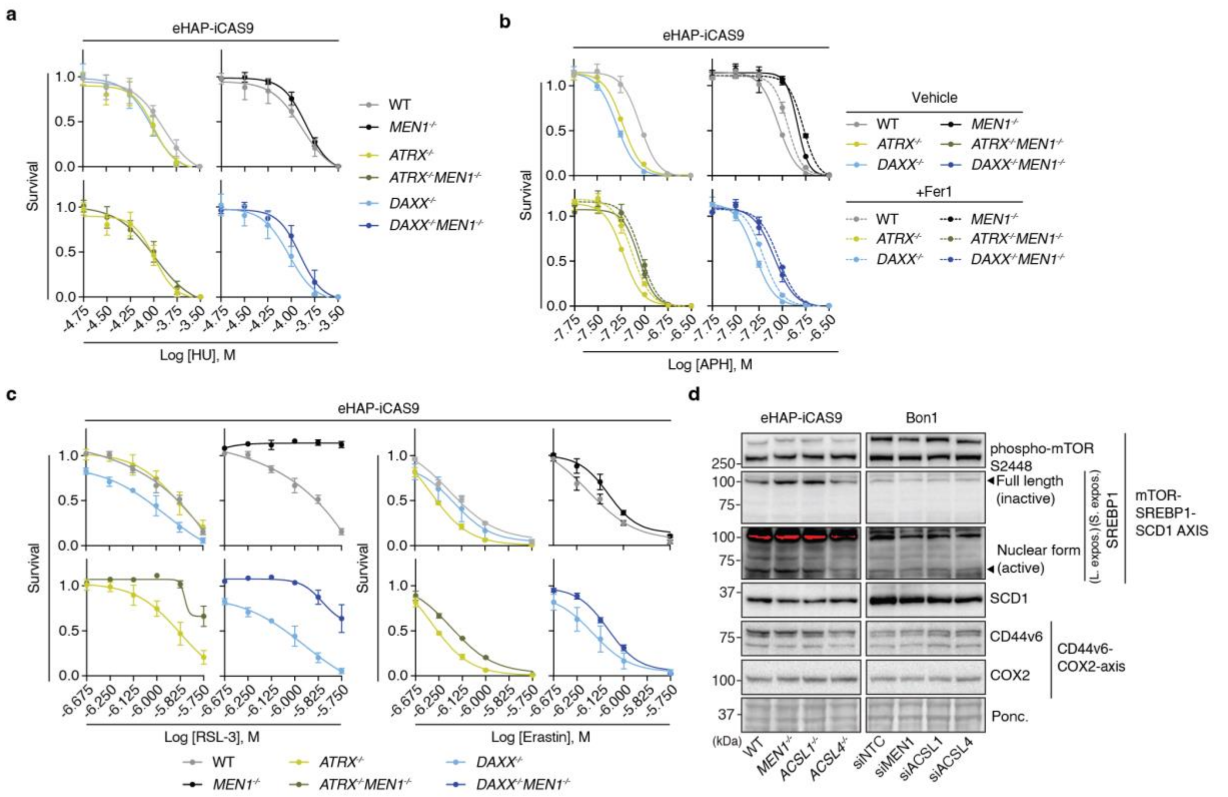


**Extended Data 2| MEN1 deficiency confers ferroptosis resistance across ATRX- and DAXX-deficient backgrounds**

**a**, Dose-response viability assays with additional replication stress-inducing agents, such as HU in WT *MEN1*$^{-/-}$, *ATRX*$^{-/-}$, *DAXX*$^{-/-}$, *MEN1*$^{-/-}$ *ATRX*$^{-/-}$, and *MEN1*$^{-/-}$ *DAXX*$^{-/-}$ CRISPR-iCas9 cells. *MEN1*$^{-/-}$ cells show resistance across multiple agents. Mean ± SEM; n =3 independent experiments.

**b**, Dose-response viability assays of Fig. 2d with replication stress-inducing agents, APH in WT *MEN1*$^{-/-}$, *ATRX*$^{-/-}$, *DAXX*$^{-/-}$, *MEN1*$^{-/-}$ *ATRX*$^{-/-}$, and *MEN1*$^{-/-}$ *DAXX*$^{-/-}$ CRISPR-iCas9 cells. Cells were also treated with ferroptosis inhibitor, Fer1, and cell death is rescued in MEN1 WT but not *MEN1*$^{-/-}$ cells in all genotypes. Mean ± SEM; n =3 independent experiments.

**c**, Dose-response viability assays of Fig. 2e with ferroptosis-inducing agents, RSL3 and erastin (Fig. 2e) in WT *MEN1*$^{-/-}$, *ATRX*$^{-/-}$, *DAXX*$^{-/-}$, *MEN1*$^{-/-}$ *ATRX*$^{-/-}$, and *MEN1*$^{-/-}$ *DAXX*$^{-/-}$ CRISPR-iCas9 cells. Mean ± SEM; n =3 independent experiments.

**d**, Immunoblot of eHAP-iCas9 cells (left; WT *MEN1*$^{-/-}$, *ACSL1*$^{-/-}$, *ACSL4*$^{-/-}$) and BON-1 cells under siRNA knockdown (right; siNTC, siMEN1, siACSL1, siACSL4), probed for proteins in mTOR-SREBP1-SCD1 axis and CD44v6-COX2 axis. Ponceau S as loading control. The protein levels are unchanged across genotypes in both cell systems.

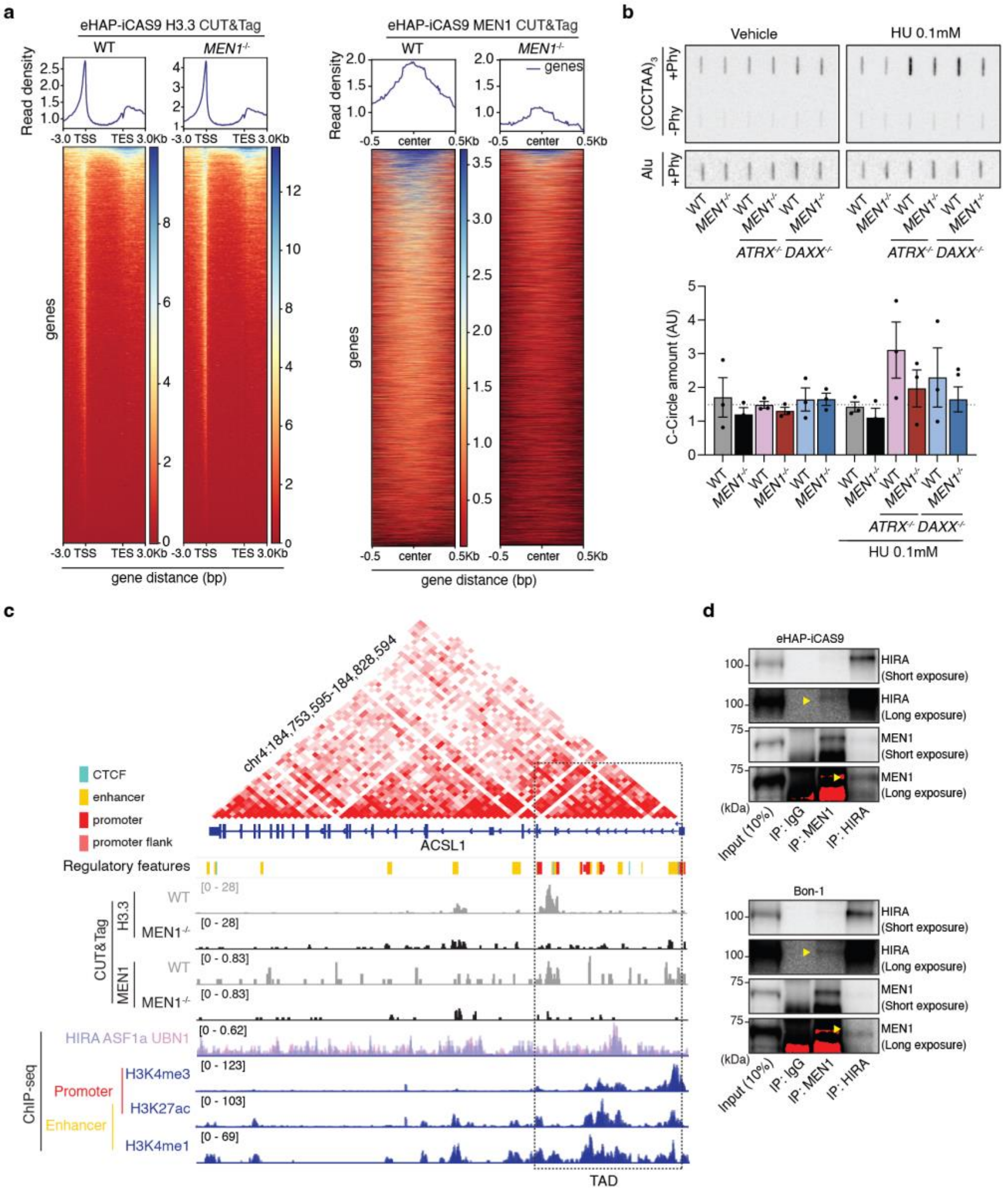


**Extended Data 3| MEN1 deficiency alters H3.3 occupancy at telomeres and ACSL1 regulatory regions**

**a**, H3.3 CUT&Tag (left) and MEN1 CUT&Tag (right) signal heatmaps in WT and *MEN1*$^{-/-}$ eHAP-iCas9 cells, centred on transcription start sites (TSS) and spanning from -3.0 kb to +3.0 kb beyond the transcription end site (TES).

**b**, Representative C-circle assay on gDNA from eHAP-iCas9 cells with indicated genotypes treated with/ without 0.1mM HU (top). Normalised C-circle levels to Alu controls (bottom). Mean ± SEM; n =3 independent samples.

**c**, Hi-C contact frequency matrix at the ACSL1 locus (chr4:184,753,595-184,828,594, GRCh38), showing the topologically associating domain (TAD) structure (dashed box)[46]. High contact frequency is shown in red; TAD boundaries are demarcated by CTCF sites. Genome browser view showing H3.3 and MEN1 CUT&Tag, and HIRA/ASF1a/UBN1 complex (GSE45024) Chip sequencing signal at the ACSL1 promoter locus[45]. Ensembl regulatory features (v115) are shown above to indicate locations of CTCF, enhancer, promoter, and promoter flank[48] and CTCF-bound elements together with active chromatin marks H3K4me3, H3K27ac, and H3K4me1[19]. MEN1, H3.3, and HIRA signals converge at the ACSL1 promoter within a single well-defined TAD, confirming three-dimensional co-occupancy at an active promoter-enhancer hub. Scale bar = 1 kb. **d**, Co-immunoprecipitation of endogenous MEN1 and HIRA in eHAP-iCas9 cells (top) and BON-1 cells (bottom). Immunoprecipitations were performed with IgG control antibody, anti-MEN1, or anti-HIRA; 10% input lysate is shown. Yellow arrowheads indicate detected bands on long exposure. It confirms a weak physical interaction between MEN1 and HIRA in both cell systems.

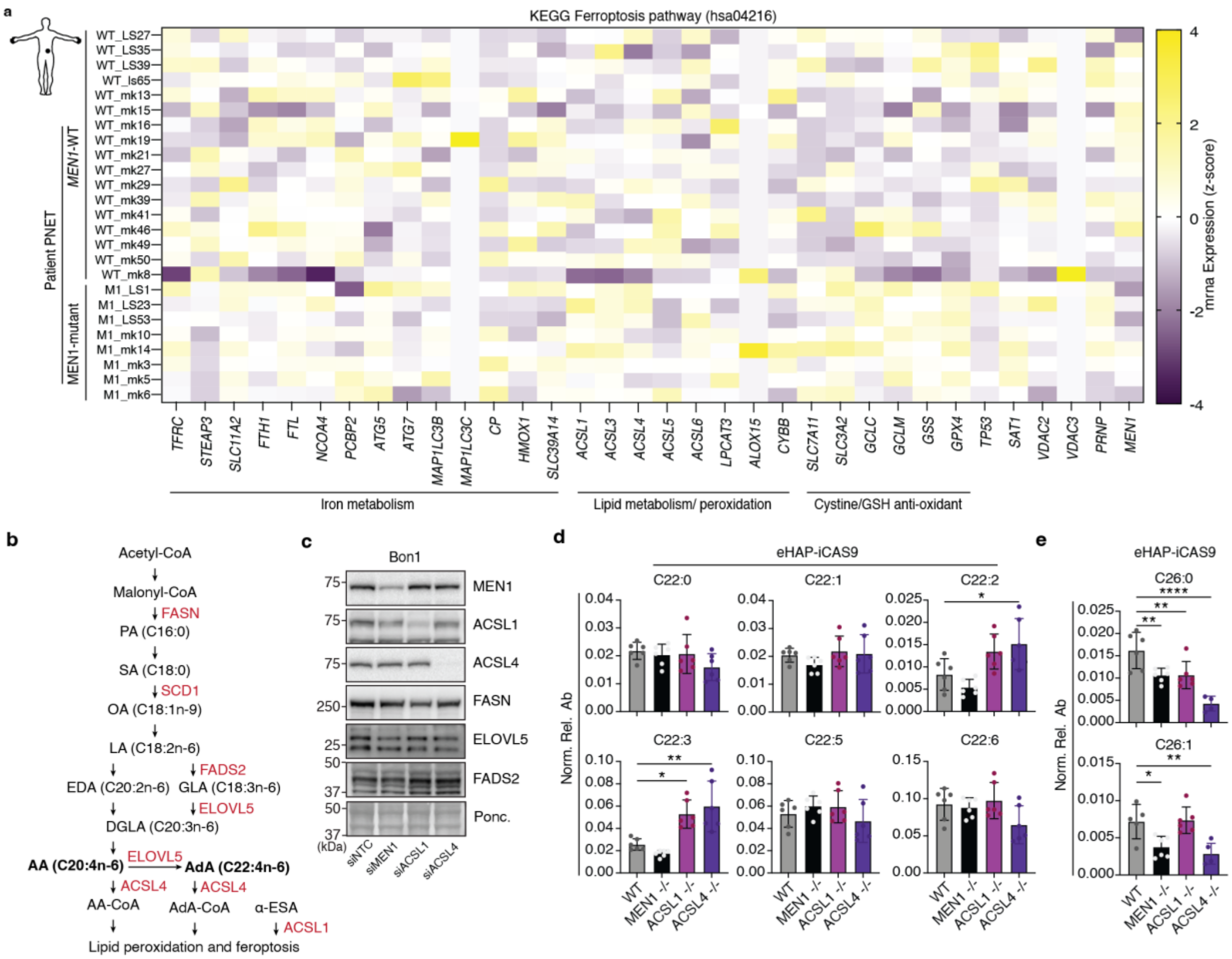


**Extended Data 4| Lipidomic and pathway analysis of MEN1- and ACSL1-deficient cells**

**a**, Ferroptosis pathway gene expression in MEN1-wild-type (n=17) and MEN1-mutant PNETs (n=8). Heatmap of expression (z-score) for 34 genes from the KEGG Ferroptosis pathway (hsa04216) across a genotyped PNET cohort[28]. Rows are grouped by functional module (iron metabolism, cystine/glutathione antioxidant metabolism, and lipid metabolism/peroxidation.

**b**, Schematic of the polyunsaturated fatty acid (PUFA) biosynthetic pathway related to lipid peroxidation and ferroptosis, annotated with enzymes relevant to this study, the pathway is adapted from[49].

**c**, Immunoblot of BON-1 PNET cells under siRNA knockdown of MEN1, ACSL1, or ACSL4 (siNTC control included), probed for proteins involved in fatty acid metabolism and ferroptosis (Extended Data Fig. 4b). Ponceau S as loading control. ELOVL5 and FADS2 are selectively reduced in siMEN1 cells in the BON-1 cells.

**d**, Dot plots of normalised relative abundance for the indicated C22 fatty acids (C22:0, C22:1, C22:2, C22:3, C22:5 and C22:6) from lipidomic profiling of WT, MEN1−/−, ACSL1−/− and ACSL4−/− eHAP-iCas9 cells. C22:2 and C22:3 are elevated in ACSL1−/− and ACSL4−/− cells (*p < 0.05, **p < 0.01; one-way ANOVA with Tukey's post hoc test). Mean ± SEM; n = 5 independent replicates.

**e**, Dot plots of normalised relative abundance for C26:0 and C26:1. Both species are significantly reduced in ACSL4−/− cells relative to WT (*$p < 0.05$, **$p < 0.01$, ****$p < 0.0001$; one-way ANOVA with Tukey's post hoc test). Mean ± SEM; n = 5 independent replicates.

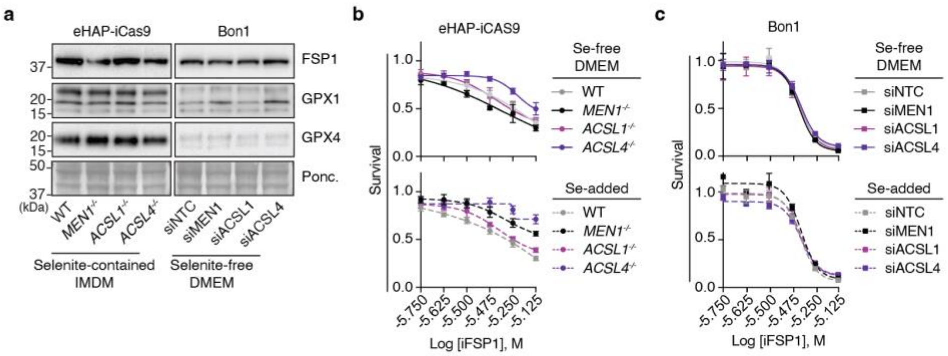


**Extended Data 5| Selenite selectively induces GPX4 and the ferroptosis phenotype is FSP1-independent**

**a**, Immunoblot showing major antioxidants that reduce lipid peroxidation in WT *MEN1*$^{-/-}$, *ACSL1*$^{-/-}$ or *ACSL4*$^{-/-}$ eHAP-iCas9 cells (left) and BON-1 cells with siRNA knockdown of MEN1, ACSL1, or ACSL4 (right). Ponceau S as loading control. All wells were loaded with equal amount of protein with the same exposure time. There is reduce expression of GPX1 and GPX4 in BON1 cells grown in DMEM in comparison to eHAP-iCas9 cells grown in IMDM.

**b,c**, iFSP1 dose-response cell viability assays in eHAP-iCas9 cells (**b**) and BON-1 cells under siRNA knockdown (**c**), cultured in selenite-free DMEM (top) or selenite-supplemented DMEM (11 ng/mL $Na_2SeO_3$, bottom). No differential response is observed between treatments with/without selenite Mean ± SEM; n =3 independent experiments.

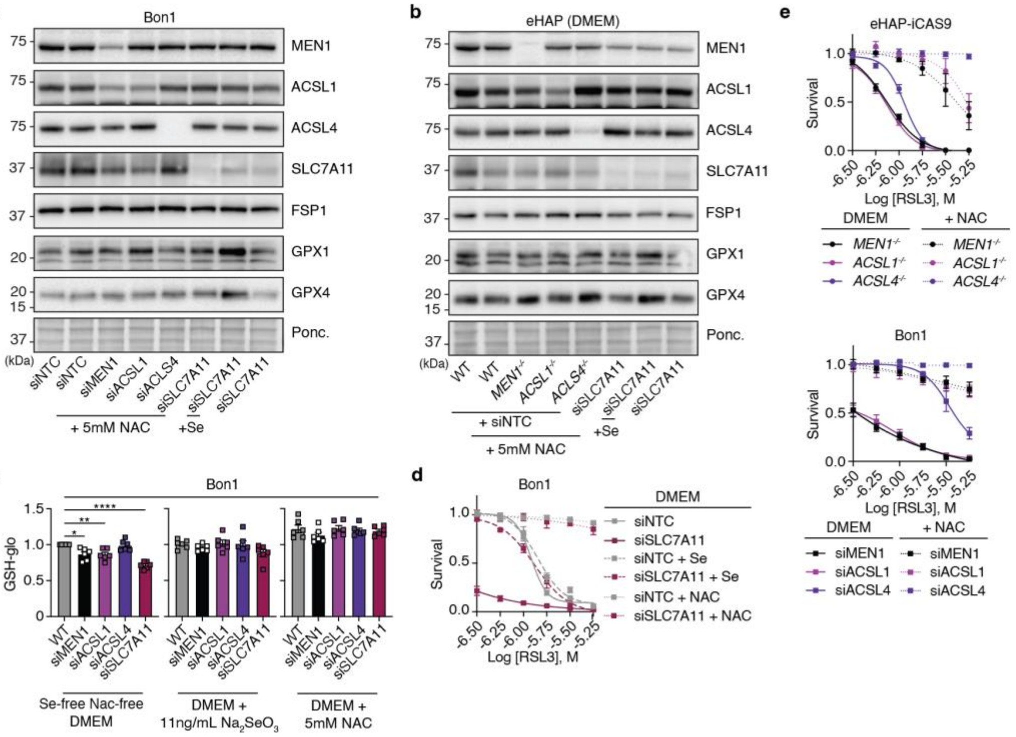


**Extended Data 6| Selenite-driven GPX4 upregulation and GSH restoration are independent of the SLC7A11-cystine axis.**

**a,b,** Immunoblots of BON-1 cells (**a**) and eHAP-iCas9 cells cultured in DMEM (**b**) under the indicated siRNA or genetic conditions ± 5 mM N-acetylcysteine (NAC) or 11 ng/mL $Na_2SeO_3$ supplementation. GPX4 is induced by selenite in siSLC7A11 and *MEN1*$^{-/-}$ conditions in both cell systems, but not by NAC supplementation, demonstrating that selenite-driven GPX4 upregulation is independent of the SLC7A11-cystine-glutathione axis.

**c**, GSH-Glo luminescent assay quantifying intracellular glutathione (GSH) in BON-1 cells of the indicated genotypes and siRNA knockdown conditions alongside siNTC controls, across three conditions: selenite-free NAC-free DMEM (left), DMEM + 11 ng/mL $Na_2SeO_3$ (centre), and DMEM + 5 mM NAC (right). Values are normalised to WT controls. GSH is significantly reduced in *MEN1*$^{-/-}$ and siSLC7A11 cells in selenite-free conditions. Selenite and NAC supplementation restore GSH levels. (*$p < 0.05$, **$p < 0.01$, ***$p < 0.001$, ****$p < 0.0001$; one-way ANOVA with Tukey's post hoc test). Mean ± SEM; n =3 independent experiments.

**d**, RSL3 dose-response cell viability assays in BON-1 cells cultured in selenite-free DMEM with indicated siRNAs treatment supplemented with 11 ng/mL $Na_2SeO_3$ or 5 mM N-acetylcysteine (NAC). siSLC7A11 cells acquire RSL3 resistance specifically upon selenite supplementation, phenocopying siMEN1 and siACSL1 conditions. NAC supplementation rescues RSL3 sensitivity. Mean ± SEM; n =3 independent experiments.

**e**, RSL3 dose-response cell viability assays in the indicated genotypes or siRNA knockdowns in eHAP-iCas9 (top) and BON-1 cells (bottom), supplemented with 5 mM N-acetylcysteine (NAC). NAC supplementation rescues RSL3 sensitivity. Mean ± SEM; n =3 independent experiments.
